# Tyrosine phosphorylation and the inhibitory C-terminal SAM domain regulate the EphA2 intracellular fragment in solution: A combined experimental and molecular modeling study

**DOI:** 10.1101/2025.09.29.679228

**Authors:** Pravesh Shrestha, Amita Rani Sahoo, Maria Iannucci, Belinda Willard, Matthias Buck

## Abstract

Eph receptors, the largest family of receptor tyrosine kinases (RTKs), orchestrate diverse developmental processes, including axon guidance, tissue patterning, and cell positioning, while also contributing to adult functions such as synaptogenesis and tissue homeostasis. Proteolytic cleavage of Eph receptors by neurodegeneration-associated proteases generates a membrane-detached intracellular region (ICR) fragment whose structural organization and signaling remain poorly understood. Here, we characterize a model for such a fragment from EphA2 and discover that the juxtamembrane region, the kinase domain, and the C-terminally linked sterile α motif (SAM) domain comprise a configurational ensemble of domain contacts and orientations which inhibit the intermolecular interactions and tyrosine cross-phosphorylation. Recombinant EphA2 ICR expressed in *E. coli* retains catalytic activity and undergoes autophosphorylation at multiple tyrosine residues which have previously been identified in eukaryotic systems. Phosphorylation weakens intramolecular interactions, as measured by microscale thermophoresis, indicating a phosphorylation-dependent remodeling of the ICR. While AlphaFold2-Multimer (AF2M) and Alphafold3 (AF3) predictions as well as available crystal structures provide only limited insight into these configurational states, coarse-grained molecular dynamics simulations using the Martini 3 force field reveal highly dynamic and predominantly transient interactions between the kinase and SAM domains that cluster at two discrete regions on the kinase surface. NMR spectroscopy of isotopically labeled EphA2 SAM suggests non-canonical intramolecular contacts that preserve the canonical SAM interaction interface for binding of partner proteins. Addition of the SHIP2 SAM domain promotes formation of the canonical EphA2–SHIP2 SAM complex while also revealing previously unrecognized interactions of the kinase domain on the SHIP2 SAM domain. Collectively, these findings establish the EphA2 ICR as a dynamic, phosphorylation-responsive signaling module in which transient intramolecular interactions regulate accessibility of several interfaces for binding. Thus, the proteolytically released EphA2 intracellular fragment is likely to have distinct characteristics in cancer and neurodegenerative disease, compared to the membrane bound form in the full receptor.

## Introduction

As cell surface receptors, receptor tyrosine kinases (<u>RTKs</u>) play critical roles as key regulators of cellular processes, such as proliferation and differentiation, tissue organization, cell migration as well as in many pathological processes (Blume-Jensen & Hunter, 2001; Lemmon & Schlessinger, 2010). With 14 members expressed in humans, Eph receptors are the largest subfamily among the RTKs (Egea & Klein, 2007). The importance of Eph/ephrin system as a multifaceted and essential regulator of developmental and disease processes is becoming well established (Kania & Klein, 2016). Studies have shown that abnormally regulated Eph participate in diverse disease processes including cataracts, neurological disorders, viral infections as well as cancer (Miao & Wang, 2012; Pasquale, 2010). In its canonical activity EphA2 signaling is mediated by ligand dependent mechanisms involving its intracellular tyrosine kinase function that regulates cell adhesion/repulsion through recruitment of adaptor and phosphorylation of partner proteins (Cioce & Fazio, 2021; Pasquale, 2024a). In addition, and in certain settings as an alternative, EphA2 functions by a ligand independent (non-canonical) signaling mechanism characterized by the phosphorylation of residues in the kinase domain-SAM domain linker, principally Ser897 (Miao et al., 2009). Thus, increased level of EphA2 expression is often associated with several advanced cancers and is linked to poor patient survival (Hess et al., 2001; Wang et al., 2008; Wykosky & Debinski, 2008).

Given its role as a critical tumorigenic and tumor-promoting factor, EphA2 has been a target for the development of cancer therapeutics (Pasquale, 2024b). More recently it has become known that EphA2 as well as other relatives are processed by proteases which are associated with neurodegenerative diseases, such as α-and γ-secretases (Merilahti & Elenius, 2019), yielding a near full-length intracellular region (<u>ICR</u>) fragment which has not yet been well characterized (Huang, 2021; Ikeda et al., 2023).

Eph receptors are unique amongst RTKs in that their ICR region possess a C-terminal sterile α motif, referred to as a SAM domain (Lee et al., 2012; Pasquale, 2024a). While SAM domains are known as binding partners for adaptor proteins and can help form homodimers, heterodimer and higher order polymer structures with other SAM domains (Borthakur et al., 2014; Knight et al., 2011), the molecular level role of the SAM domain for EphA2 function is only revealed gradually through studies over the last several years. Specifically, work by us and others has shown that the presence of the EphA2 SAM domain appears to inhibit kinase activity in cells and, conversely, exhibit increased oligomerization when the SAM domain deleted (Lechtenberg et al., 2021; Shi et al., 2017; Singh et al., 2017). Recently, Pasquale and colleagues published the x-Ray crystal structure of an EphA2 intracellular region (albeit with an N-terminally truncated juxtamembrane segment) (Lechtenberg et al., 2021). However, it is suspected that the Kinase-SAM domain interactions may only exist in the context of the crystal lattice. Furthermore, a significant change in the biophysical behavior and in the crystal structure was seen with 5 phosphomimetic mutations in the linker surrounding Ser 897, suggesting that multiple Ser/Thr linker phosphorylation events can cooperatively regulate the conformational landscape of EphA2 ICR (Lechtenberg et al., 2021). (This topic is being investigated further in our lab and will be subject of a separate report).

Here, we examined the potential relationships between the SAM domain, EphA2 kinase activity, auto-and trans-phosphorylation, and ICR dimerization/oligomerization using purified EphA2 ICR, and a SAM domain-deleted ICR fragment, ICRΔSAM in solution under controlled conditions. We focus on ICR using two important EphA2 models: the full-length here after referred to as ICR and the ICR which is truncated in the linker and missing its SAM domain as ICRΔSAM. We investigated the role of the EphA2 SAM domain and ICR tyrosine phosphorylation on the dimerization and activity of these two models, as well as on their dephosphorylated forms (either phosphatase treated or a kinase dead mutant). Results from microscale thermophoresis (MST) using the ICR and ICRΔSAM proteins support an inhibitory role for the EphA2 SAM domain in receptor dimerization in solution. These observations are elaborated by molecular modeling using Coarse Grained (Martini 3.0) dimerization simulations, but also by simulations in which SAM domain containing fragments are added to the ICR/ICRΔSAM. NMR titrations together with MST further confirmed the binding of an additional EphA2 SAM domain to the ICR, suggesting that the interaction involves contacts outside the canonical contacts of the EphA2 SAM domain.

The EphA2 SAM domain by itself does not dimerize (Borthakur et al., 2014; Lee et al., 2012; Shi et al., 2017), it interacts with the SAM domain of SHIP2, SH2 domain-containing inositol 5-phosphatase 2 (an intracellular enzyme that regulates cell signaling by dephosphorylating the lipid second messenger PI(3,4,5)P₃ into PI(3,4)P₂) to form a heterotypic, albeit dynamic SAM–SAM complex that regulates EphA2 signaling, trafficking, and cellular functions (Zhang et al., 2016). Thus this interaction has emerged as a potential target for therapeutic intervention (Lee et al., 2012; Leone et al., 2008; Wang et al., 2018; Zhuang et al., 2007).

Overall, our findings point to new interfaces and key residues which can now be mutated for further in vitro and in-cell studies but they also suggest that domain-domain interactions are transient and may become more defined as higher order interactions, such as those with the plasma membrane, and domains in the full-length receptor are added.

## Results

### EphA2 Intracellular region purified from *E. coli* is phosphorylated and active

The human EphA2 intracellular region (ICR; residues 559–976) was expressed in *E. coli* with an N-terminal TRX–His tag containing a TEV protease site (Fig. 1A) as done in two previous reports by another lab (Zabell et al., 2006;Blaasubramaniam et al., 2011), but with a slightly modified protocol (see supplementary methods). The TRX tag improved expression and folding, while the His tag enabled affinity purification ( Zabell et al., 2006; Gande et al., 2016). The tag was removed by TEV cleavage (Fig. S1A), yielding pure wildtype (wt) or kinase-dead (D739N) ICR (∼50 kDa) and ICRΔSAM (∼40 kDa), which migrated as monomers on Superdex 75 size-exclusion chromatography (Fig. S1B). Western blotting with intracellular region (550-600) and pY594-specific antibody showed tyrosine phosphorylation of wt ICR and ICRΔSAM but not the kinase-dead variant (Fig. 1B), consistent with the absence of endogenous tyrosine kinase activity in *E. coli*. LC-MS analysis confirmed no phosphorylation in the kinase dead (data not shown) but phosphorylation at Y588, Y594 in the juxtamembrane region and Y628, Y772 in the both wt ICR and ICRΔSAM, residues critical for EphA2 activity (Balasubramaniam et al., 2011; Wiesner et al., 2006). However, phosphorylation at SAM-domain tyrosines (Y921, Y930, and Y960) was not detected. The difference in SAM-domain phosphorylation may reflect variations in experimental conditions, protein preparation, phosphorylation efficiency, local residue accessibility, or limitations in mass spectrometric detection. Importantly, the lack of detected phosphorylation at SAM tyrosines does not contradict previous findings but highlights the potential heterogeneity of EphA2 phosphorylation patterns under different experimental contexts. We see a relatively modest difference in phosphorylation between ICR and ICRΔSAM (Fig. 1C). Data are deposited under ProteomeXchange PXD068904.

**Figure 1.**
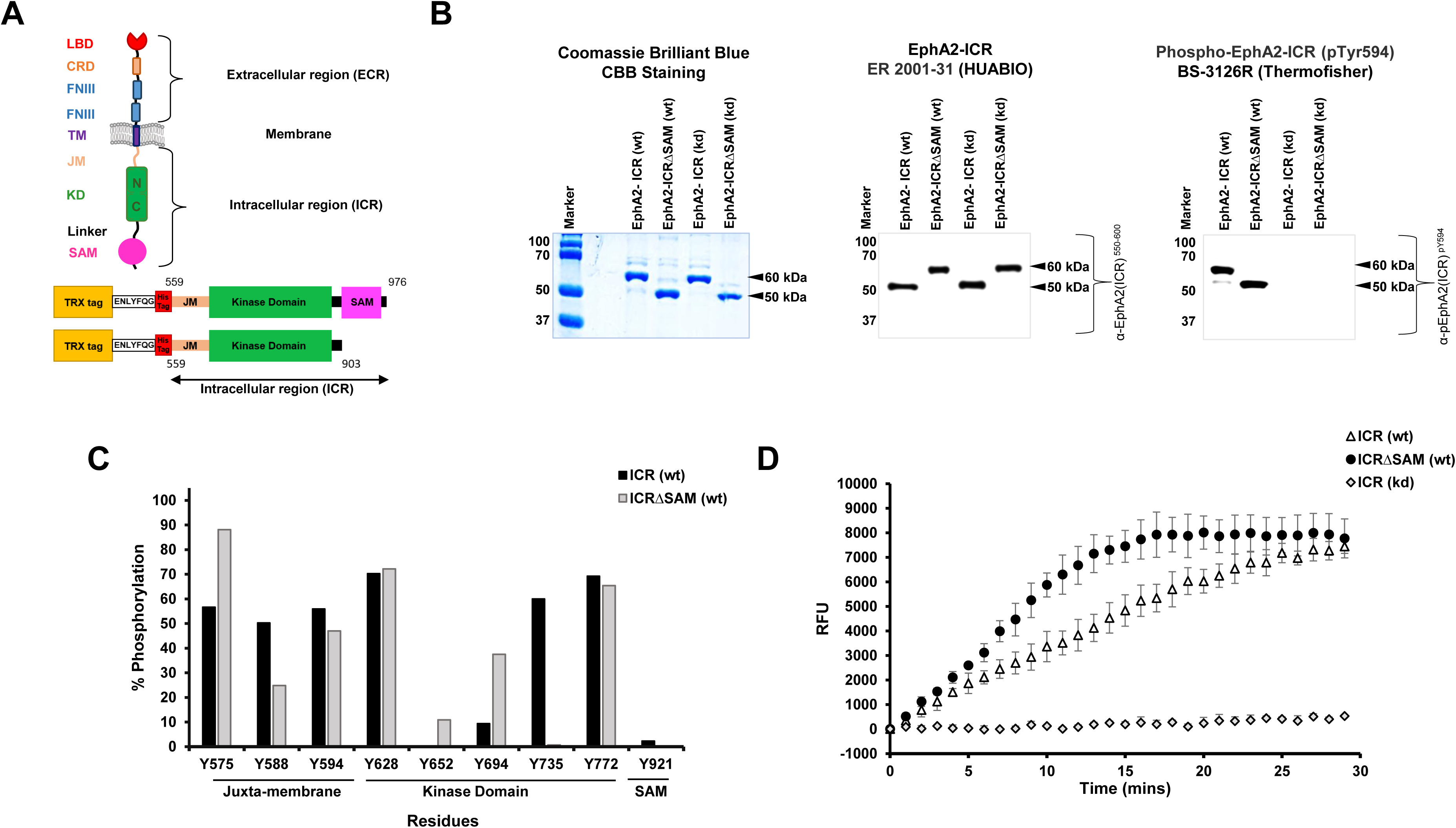
Cloning, purification and characterization of the EphA2 intracellular region (ICR, ICRΔSAM) (A) Domain architecture and construct design. (B) Purified protein detection using coomassie brilliant blue, CBB staining (far left), antibody against intracellular region (550-600), HUABIO (middle panel) and Juxta-membrane specific phospho-tyrosine (pY594), Thermofisher (far right panel). (C) Comparison of residue specific phosphorylation of both wt ICR and wt ICRΔSAM using LC-MS analysis. (D) PhosphoSens-Kinetic kinase activity assay of wt ICR, ICRΔSAM and kd ICR using synthetic phosphoSens kinase substrate.

PhosphoSens kinase activity assays using an EphA2-specific synthetic peptide substrate revealed that the ICR exhibited an average 44% reduction in kinase activity compared with the ICRΔSAM construct, with initial reaction rates of 329 ± 50 RFU/min and 584 ± 70 RFU/min, respectively (Fig. 1D). The faster initial reaction rate of ICRΔSAM likely reflects the cumulative effect of phosphorylation and/or configurational changes associated with the deletion of the SAM domain, such as the increased frequency of forming transient intermolecular dimers (see below), rather than large differences at individual phosphorylation sites. These results suggest that the C-terminal SAM domain exerts an inhibitory regulatory effect on EphA2 kinase activity, potentially by modulating the configurational dynamics or accessibility of the catalytic domain.

### EphA2 SAM domain inhibits the dimerization/oligomerization of the EphA2 intracellular domain

Ligand-dependent dimerization/oligomerization and ligand-independent phosphorylation of full-length EphA2 via changes in its extracellular region are well established (Shi et al., 2023; Singh et al., 2018; Xu et al., 2011). However, the residue-level mechanisms governing intracellular region (ICR) dimerization or oligomerization, particularly in the absence of ligand or extracellular/transmembrane domains, remain unclear (Pasquale, 2024a; Sahoo & Buck, 2021; Zapata-Mercado et al., 2025). Here, we examined ICR self-association using MST. Wildtype ICR showed modest binding with a dissociation constant (Kd) of 15.0 ± 5.0 µM (Fig. 2A). Deletion of the SAM domain significantly enhanced binding, yielding a Kd of 5.4 ± 0.5 µM (Fig. 2B). To assess the role of phosphorylation, kinase-dead ICR and ICRΔSAM were analyzed and displayed stronger interactions, with Kd values of 1.3 ± 0.1 µM and 1.0 ± 0.1 µM (Fig. 2C and 2D) respectively. These results indicate that SAM domain removal and loss of phosphorylation similarly increase ICR affinity, although SAM deletion has a much smaller effect in the non-phosphorylated state. Since MSTs sensitivity to transient interactions differs from the gel filtration results, we also used an alternative fluorescent label give approximately the same result (see supplemental methods and discussion therein). Self-association of all ICR variants was further confirmed by chemical cross-linking assays using commercially available cross linkers (DSSO and DC4) for both wt ICR and ICRΔSAM (Suppl. Information Fig. S1C) and kinase dead ICR and ICRΔSAM (Fig. S1D), demonstrating formation of dimers and higher-order oligomers with considerable dependence on phosphorylation or kinase activity.

**Figure 2.**
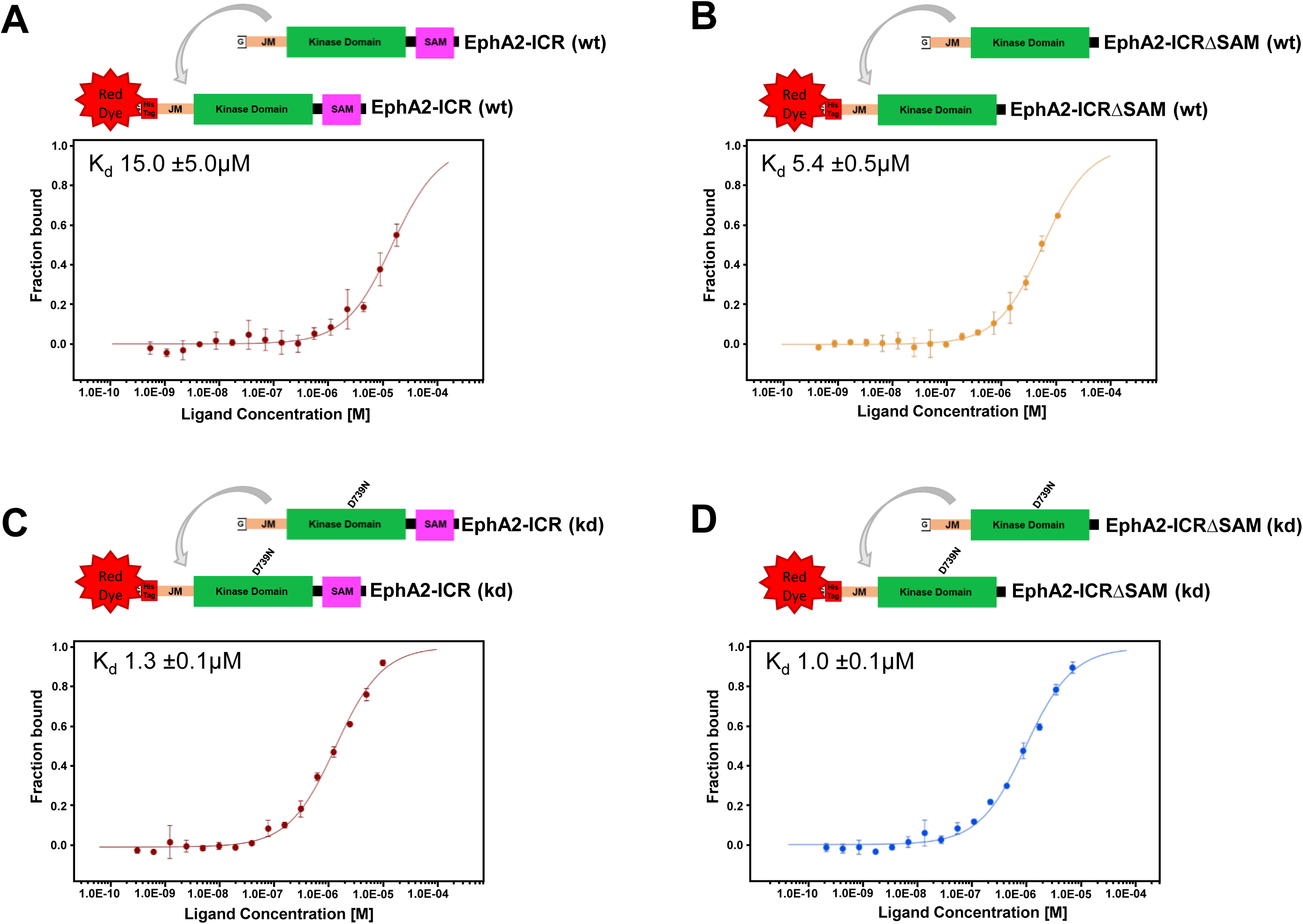
Phosphorylation regulates the association of EphA2 ICR, ICRΔSAM. (A) Association between wt ICR; (B) wt ICRΔSAM; (C) kd ICR and (D) kd ICRΔSAM using MST.

### Coarse grained MD simulations suggest models for monomers and dimers which are more consistent than X-ray crystal structures or Alphafold predictions

For each system (ICR monomer, ICR dimer, and ICRΔSAM dimer) we performed four independent 4 μs coarse-grained (CG) molecular dynamics simulations. Dimer simulations were initiated from unphosphorylated monomers placed 5.0 nm apart with randomized orientations and velocities. Structural ensembles were analyzed by RMSD-based clustering using cutoffs of 0.6 nm for monomers and 0.8 nm for dimers.

ICR monomer simulations yielded two dominant structural clusters, comprising ∼40% and ∼30% of the population, distinguished by different relative orientations of the SAM and kinase domains (Fig. 3A). In the most populated configuration, the SAM domain interacts extensively with the juxtamembrane (JM) region and the N-lobe of the kinase domain, adopting an “upward” orientation that suggests a potential autoinhibitory role by restricting kinase accessibility. In the second cluster, the SAM domain associates with the lower face of the kinase C-lobe. These results highlight substantial interdomain flexibility and a dynamic SAM–kinase interface (Table S1) even in the absence of intermolecular interactions, such as those seen in a dimer.

**Figure 3.**
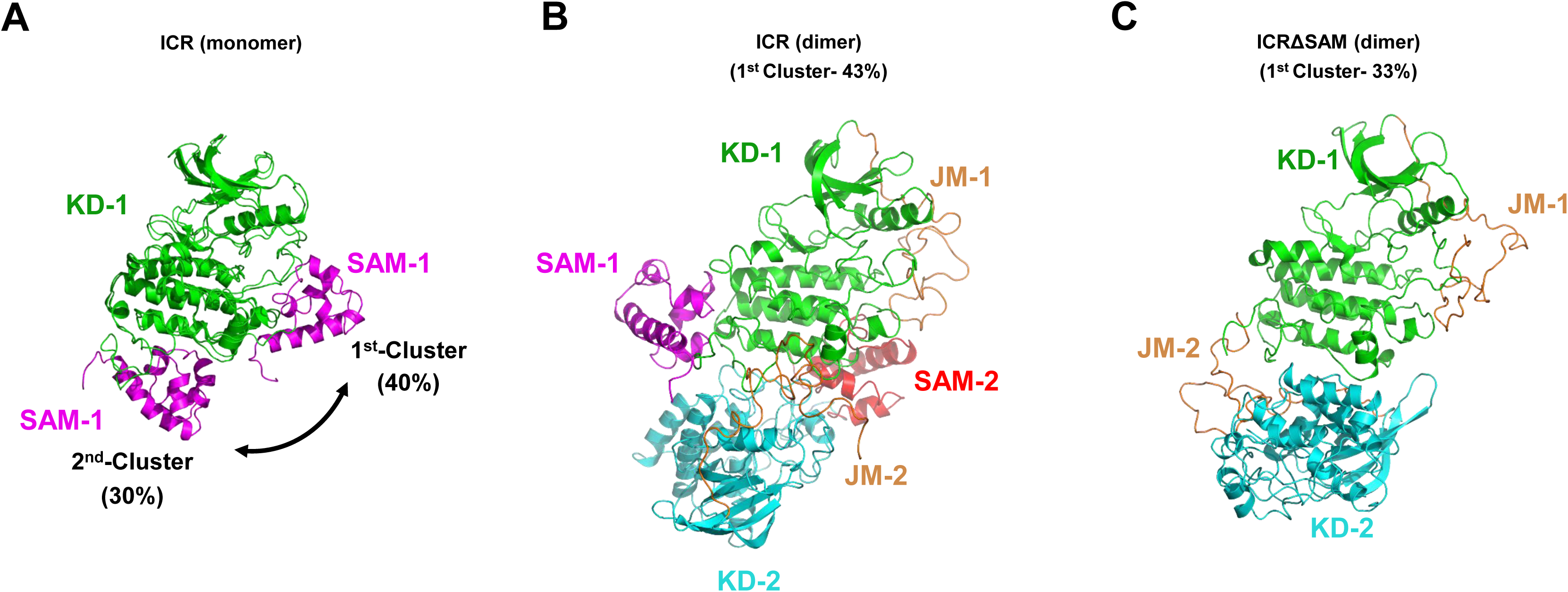
Overall comparison of wt ICR monomer, ICR dimer and ICRΔSAM dimer configurations obtained from CG simulation. (A) Structural alignment of the central structures of top two clusters of ICR monomer showing the possible orientation of the SAM domain (magenta) with respect to the kinase domain (green). The central structures from the top cluster of the wt ICR dimer (B) and ICRΔSAM dimer (C) obtained from CG simulations. JM regions of both chains are shown in orange, respective kinase domains (green-chain A; cyan-chain B) and respective SAM domains (magenta-chain A; red-chain B).

In ICR dimer simulations, the most populated cluster (43%) revealed a compact, asymmetric dimer stabilized primarily by KD-SAM and KD-KD interactions, with additional contacts involving one JM region (Fig. 3B). Persistent interactions between SAM and kinase domains further stabilized the dimer architecture, yielding an ensemble of compact conformations that persisted across simulations and are consistent with a functional signaling assembly (Fig. S3).

In contrast, the ICRΔSAM dimer adopted a distinct organization. The dominant cluster (33%) showed altered positioning of the JM-2 and KD-2 domains when aligned to the ICR dimer, reflecting the loss of SAM-mediated contacts that otherwise help orient the kinase domains (Fig. 3C and S2B). Contact maps confirmed differences in KD and JM interactions between ICR and ICRΔSAM dimers (Fig. S4). Notably, both constructs consistently favored asymmetric dimer topologies.

Comparison with available EphA2 crystal structures, including a near full-length ICR dimer (Lechtenberg et al., 2021), revealed similarities in overall asymmetry but fewer inter-protein contacts in the crystal structures than in the simulation-derived dimers (Fig. S5A; Table S2), suggesting that the CG models may capture more extensive solution-state interactions. Previous crystal structures of truncated EphA2 constructs exhibit additional dimer interfaces that have been interpreted as crystal lattice contacts in the original structural studies. Because these interfaces differ substantially among crystal forms and involve truncated constructs lacking either the JM or SAM domains (Xu et al., 2015), we consider them less likely to represent the predominant solution-state assembly studied here (Fig. S5B, Table S3).

To assess structural tendencies of these dimers and evaluate predictive modeling, dimeric models of ICR and ICRΔSAM were generated using AF2M and AF3 (Fig. S6, Table S4). AF2M ICRΔSAM dimer model showed higher confidence (>0.60 fair-to-good threshold), the full-length ICR prediction showed substantially lower confidence, suggesting that the presence of flexible linker and SAM regions reduces prediction reliability. Both models predicted symmetric dimers, with side-by-side SAM– SAM interfaces for ICR and head-to-head N-lobe kinase interactions for ICRΔSAM. Consistent with the ICR crystal structure (PDB: 7KJA), AF2M predicts SAM contacting the KD C-lobe via helix 5, whereas CG simulations reveal an alternative interface involving the SAM middle linker, suggesting context-dependent SAM reorientation between monomeric and dimeric states. Moreover, both the ICR and ICRΔSAM AF3 models exhibited very low interface confidence scores, indicating considerable uncertainty in the predicted interfaces. Interestingly, despite these low confidence values, the relative orientation of the kinase domains in the AF3 ICR model resembles that observed in the CG simulations. Taken together, these results suggest that while AlphaFold predictions provide useful starting models, they currently have limited ability to resolve the dynamic multidomain organization of the EphA2 intracellular region. Although AF2M predicted relatively confident symmetric dimers for ICRΔSAM, neither AF2M nor AF3 reproduced the asymmetric donor–acceptor kinase dimer topology observed in experimentally determined Eph receptor and EGFR kinase dimers. In contrast, the dominant CG-derived dimers consistently adopted asymmetric arrangements for both ICR and ICRΔSAM. While this observation does not establish the CG models as uniquely correct, it indicates that the CG simulations better reproduce experimentally established kinase dimer architectures than the AlphaFold-derived models.

### EphA2 binding to domains: Inter-protein SAM interactions modestly relieve autoinhibition

Having established that the SAM domain modestly influences ICR dimerization, we next tested whether addition of isolated SAM domains, either EphA2 SAM or its binding partner SHIP2 SAM, could compete with kinase–SAM interactions and enhance kinase activity. Time-dependent transphosphorylation assays were performed using wildtype (wt) ICR or wt ICRΔSAM as enzymes and kinase-dead (kd) ICR as an EphA2-specific substrate (Fig. 4). The kd substrate retained its TRX tag, increasing its size (∼60 kDa) and allowing it to be clearly distinguished from the active kinase on the gel. With wt ICR as the enzyme, phosphorylation of kd ICR at Tyr594 was detectable at 17% (scaled to most intense, here final time-point) after 10mins, around 55% at 30 mins and complete by 90 min after ATP addition (Fig. 4A left upper panel). In contrast, wt ICRΔSAM phosphorylated the substrate to 45% after 10 min and almost completely at 85% after 30 mins (Fig. 4B right upper panel), confirming the inhibitory role of the EphA2 SAM domain. Addition of isolated EphA2 SAM had no visible effect on wt ICR activity, with comparable substrate phosphorylation of ICR only (Fig. 4C middle left panel). Conversely, the addition of an EphA2 SAM domain in trans slightly reduced the activity of wt ICRΔSAM, as indicated by decreased phosphorylation at early time points (Fig. 4D middle right panel). MST detected no binding of EphA2 SAM to wt ICR or ICRΔSAM but revealed modest binding to kd ICR and kd ICRΔSAM (data not shown), indicating the presence of additional SAM-interacting sites within the ICR that can modulate intermolecular interactions and transphosphorylation.

**Figure 4.**
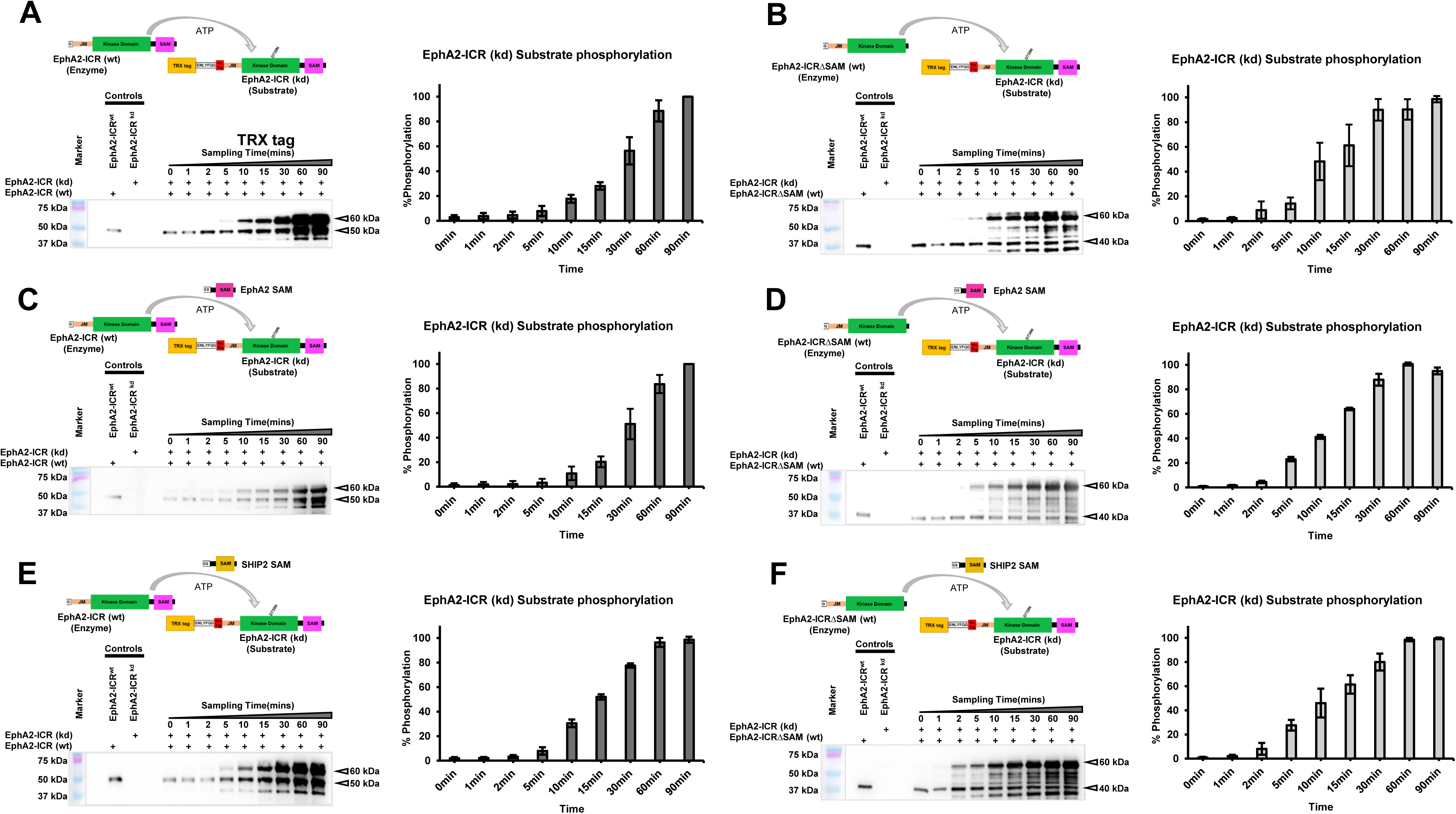
EphA2 SAM and SHIP2 SAM domain modulate the kinase activity of intracellular region of EphA2. Time dependent transphosphorylation of kd EphA2 ICR by (A) wt ICR and (B) wt ICRΔSAM using in-vitro transphosphorylation assay in presence of EphA2 SAM domain (C and D) and in presence of SHIP2 SAM domain ( E and F). 10µM ICR kd (Substrate) was incubated with 250nM ICR and 250nM ICRΔSAM in presence of 1mM ATP and in presence of 10µM SAM domain of EphA2 and SHIP2. Visualizations of western blots using antibody developed against pY594 EphA2, thermofisher. Right panels show normalized quantification of each blot using loading controls. Quantification of the blots were performed using Image J software.

Addition of the SHIP2 SAM domain to wt ICR accelerated ICR phosphorylation, reaching approximately 30% phosphorylation after 10 min, ∼75% after 30 min, and near completion by 60 min (Fig. 4E, lower left panel). In contrast, no enhancement of phosphorylation was observed with wt ICRΔSAM (Fig. 4F, lower right panel), likely due to the absence of the EphA2 SAM domain required for the SHIP2 SAM-mediated interaction in this construct. MST further confirmed binding of SHIP2 SAM to ICR constructs, with stronger affinity for full-length ICR containing the native SAM domain (Fig. S7). Overall, these data suggest that addition of both EphA2 SAM and SHIP2 SAM domains, as “extra” proteins in trans, can relieve inhibitory SAM-mediated interactions in the full length ICR.

### Mapping the interaction between SAM domains and the ICR and ICR**Δ**SAM using solution NMR

Next, we used solution NMR to monitor the binding of EphA2 SAM to the 4 ICR constructs. The interaction between these proteins was confirmed by collecting 2D [^1^H-^15^N]-TROSY HSQC experiment of ^15^N-uniformly labeled kd ICR in presence and absence of unlabeled EphA2 SAM. Several peaks showed chemical shift perturbation as well as broadening in the TROSY spectra (Fig. S8). However, the spectra of the EphA2 ICR have not been assigned yet and the ICR proteins are difficult to handle for longer periods of time at the high concentrations (> 300 µM) needed for such NMR studies. Thus, we could not map the interface region on the side of the ICR. Instead, we used the spectra of the EphA2 and SHIP2 SAM domains, assigned previously (Lee et al., 2012), as a read-out of interactions the added SAM domains make with the ICR and ICRΔSAM in both phosphorylated (wt) as well as dephosphorylated/kd forms. Importantly, we had shown previously that neither EphA2 SAM nor SHIP2 SAM domains self-dimerize in solution (Lee et al., 2012), the former being relevant here, as SAM domains of at least some Eph receptors are used to dimerize/polymerize them (Singh et al., 2015).

First, 2D [^1^H-^15^N]-HSQC experiments were recorded for a uniformly ^15^N labeled EphA2 SAM in absence and presence of unlabeled dephosphorylated ICR (Fig. 5A). The binding site on EphA2 SAM was mapped by comparing the peak intensity changes caused by the addition of unlabeled and dephosphorylated ICR (Fig. 5B). Except for two residues left from the N-terminal tag after cleavage, decrease in peak intensity, (ratio < 0.88) of EphA2 SAM residues map to the α2 helix of the EphA2 SAM, which runs from residues 919-928 (Fig. 5C). Remarkably, titrating phosphorylated, i.e. wt ICR and ICRΔSAM onto ^15^N labeled EphA2 SAM yielded only small changes to the peak intensities of a few EphA2 SAM signals, indicating that the binding of EphA2 SAM to phosphorylated ICR is much weaker (data not shown, but consistent with the absence of binding seen by MST). Taken together, our data from MST and the solution NMR titrations revealed regions on the added SAM domain that are involved in the molecular interaction with the ICR, in case of ICRΔSAM obviously outside of its own, linked SAM domain.

**Figure 5.**
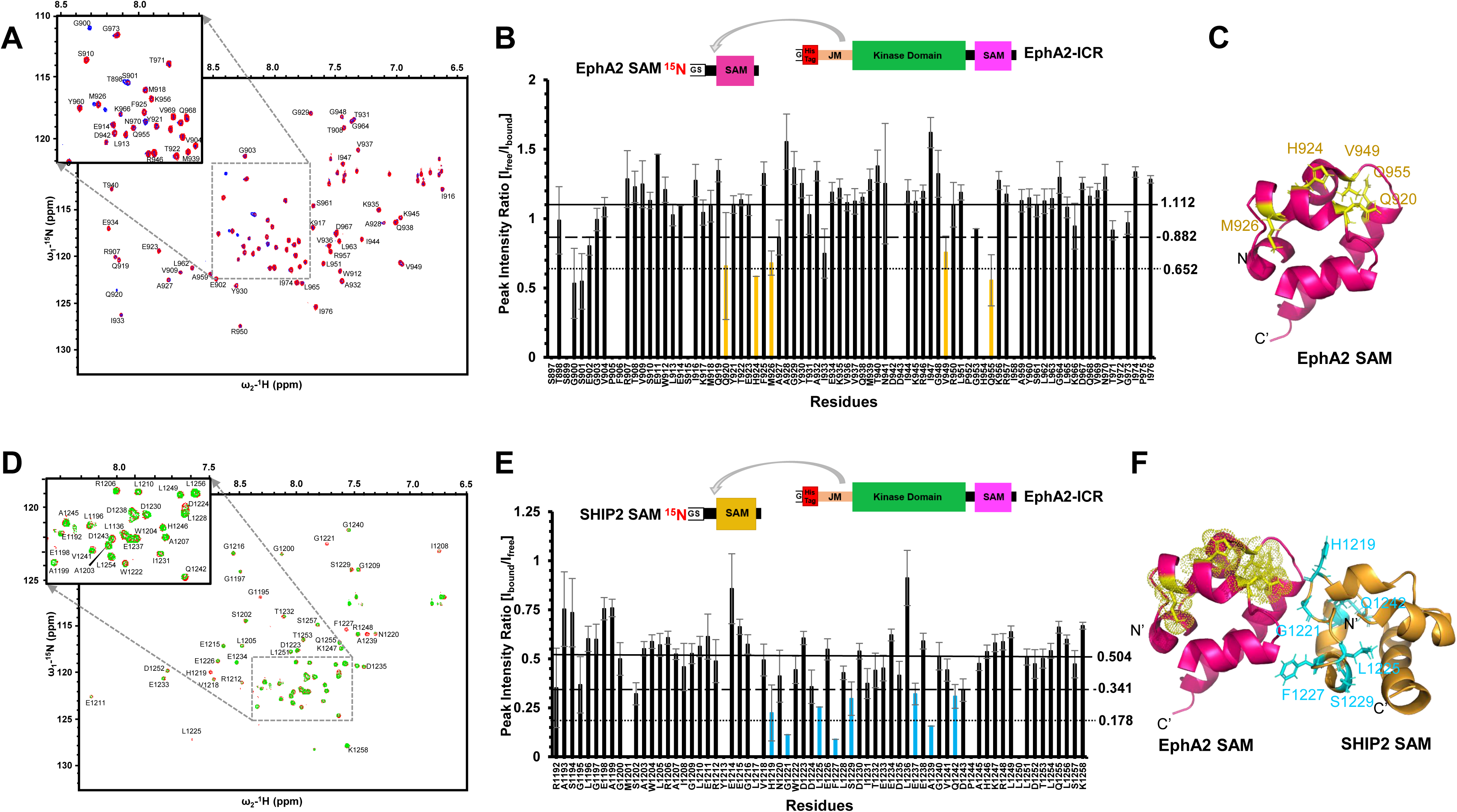
Mapping the interaction between dephosphorylated ICR and EphA2/SHIP2 SAM domain using solution NMR. (A) ^1^H-^15^N HSQC of EphA2 SAM (Red) titrated with unlabeled unphosphorylated ICR (Blue) with 1:0.5 molar ratio in 20mM TRIS pH 7.2; 150mM NaCl, 5mM MgCl_2_ & 2mM TCEP at 298K at 900MHz. (B) The peak intensity ratio of 15N labeled EphA2 SAM in the absence and presence of unlabeled EphA2 ICR (in 1:0.5 molar ratio) plotted by residue number. The upper solid and middle dashed lines represent “mean value (1.112)” and “mean value - standard deviation (0.882)” while the bottom dotted line signifies “mean value-2xstandard deviation (0.652)”. (C) EphA2 SAM domain (magenta, PDB: 2kso) with the perturbed residues mapped. (D) SHIP2 SAM interacts with ICR via the canonical mid loop region ^1^H-^15^N HSQC of SHIP2 SAM (Red) titrated with unlabeled unphosphorylated ICR (Green) at 1:0.125 molar ratio in 20mM TRIS pH 7.2; 150mM NaCl, 5mM MgCl_2_ & 2mM TCEP at 298K at 800MHz. (E) The peak intensity ratio of 15N labeled SHIP2 SAM in the absence and presence of 1:0.125 molar ratio of unlabeled EphA2 ICR was plotted against the residue numbers. The upper solid and middle dashed lines represent “mean value (0.504)” and “mean value - standard deviation (0.341)” while the bottom dotted line signifies “mean value-2 x standard deviation (0.178)”. (F) SHIP2 SAM domain (orange) with the perturbed residues mapped upon ICR interaction. EphA2 SAM (magenta) indicating the peak intensity change upon ICR interaction (yellow). PDB: 2kso

### SHIP2 SAM domain interacts with EphA2-ICR via canonical mid-loop region

We next examined interactions between SHIP2 SAM and EphA2 ICR or ICRΔSAM using MST and solution NMR. MST measurements showed that wildtype (wt) ICR binds the trans-added SHIP2 SAM with a dissociation constant (Kd) of 2.6 ± 0.9 µM (Fig. S7A), comparable to the previously reported affinity between isolated EphA2 SAM and SHIP2 SAM (Kd = 5.2 ± 1.2 µM) (Lee et al., 2012). The modest difference likely reflects the presence of the full ICR versus the isolated SAM domain and/or methodological differences between this and the previous measurements (e.g. previous measurements were done with SAM domains in presence of their His_6_-tags).

To map the interaction, 2D [^1^H–^15^N]-HSQC spectra of uniformly ^15^N-labeled SHIP2 SAM were recorded in the absence and presence of unlabeled ICR (Fig. 5D). Residues showing strong intensity reductions, ratio < 0.34 (Fig. 5E) were mapped onto the known EphA2 SAM–SHIP2 SAM heterodimer structure, PDB 2KSO (Fig. 5F), confirming engagement of the canonical SAM–SAM interface. Extensive peak broadening, consistent with intermediate exchange on the NMR timescale, indicates a low micromolar affinity interaction and formation of a larger complex. Titration of phosphorylated wt ICR and wt ICRΔSAM into ^15^N-labeled SHIP2 SAM also produced significant chemical shift perturbations (>0.025 ppm) (Fig. S9A, B). Affected residues localized primarily to the mid-loop (ML) interface and neighboring regions, again consistent with a canonical SAM–SAM interaction. There are some perturbations in other regions, in terms of chemical shift changes, which suggests that SHIP2 SAM likely also interact with the JM region, kinase domain, or linkers of the ICR. In particular, similar –albeit slightly weaker-perturbations were observed upon titration with ICRΔSAM, supporting the presence of additional SAM-binding sites within the ICR even in the absence of the as the EphA2 SAM domain primary binding site for SHIP2 SAM.

Consistent with the NMR data, phosphorylation and deletion of the SAM domain in EphA2 reduced but did not abolish SHIP2 SAM binding, with Kd values of 2.6 ± 0.9 µM for wt ICR (Fig. S7A) and 7.5 ± 2.1 µM for wt ICRΔSAM (Fig. S7B). While most affected residues overlapped between ICR and ICRΔSAM titrations, differences relative to dephosphorylated/kd ICR/ICRΔSAM, which has a significantly stronger binding suggest multiple binding modes and subtly shifted interaction surfaces on SHIP2 SAM. Overall, MST and NMR indicate that EphA2 SAM/EphA2 ICR (Fig. 5F, Yellow sphere) and SHIP2 SAM/EphA2 ICR complexes (Fig. 5F, cyan) engage for their main interactions in largely distinct interfaces on the SAM domains, highlighting the presence of multiple interaction sites in the ICR.

### EphA2 and SHIP2 SAM domains in CG Simulations and in AF2M / AF3 predictions

To determine whether simulations and structure predictions provide insights into interactions of ICR or ICRΔSAM with added EphA2 or SHIP2 SAM domains, we performed coarse-grained (CG) simulations, AF2M and AF3 predictions (Fig. S10). In all approaches, the added (non-linked) EphA2 SAM localized to a similar kinase-proximal region in the ICR (Fig. S10A, E, and I), although its orientation differed among the models. This region is also sampled by the native SAM domain in the CG simulations of the ICR monomer and dimer (Fig. 3A; Tables S5–S7). Similarly, the added EphA2 SAM bound to a comparable interface on ICRΔSAM across CG, AF2M, and AF3 models (Fig. S10B, F, and J). In contrast, greater differences were observed for SHIP2 SAM. In the full-length ICR, AF2M and AF3 predicted the canonical end-helix/mid-loop interaction between SHIP2 SAM and the covalently linked EphA2 SAM (Fig. S10G and K), whereas the CG simulations sampled an alternative arrangement in which SHIP2 SAM contacted both the EphA2 SAM and kinase domain (Fig. S10C; Tables S5–S7). For ICRΔSAM, AF2M and AF3 predicted similar SHIP2 SAM binding interfaces, whereas the CG model showed a different interface (Fig. S10D, H, and L) resembling that sampled in the CG ICR–SHIP2 SAM complex (Fig. S10C). These differences highlight the variability among the predicted interaction modes, particularly for SHIP2 SAM. Notably, the AF2M and AF3 models also differ in their interface confidence scores, which should be considered when interpreting these alternative binding modes. These CG simulation results agree with the NMR data showing chemical shift changes and line broadening in SHIP2 SAM upon ICR binding, which are reduced with ICRΔSAM. For ICRΔSAM, SHIP2 SAM shifts toward the kinase N-lobe in both AF2M and CG simulations (Fig. S10D and H). As an aside, and consistent with the experimental observations (Lee et al., 2012), AF2M predicts the EphA2– SHIP2 SAM heterodimer with high reliability, unlike the EphA2 SAM homodimer (Fig. S11).

## Discussion

Our combined biophysical and computational analyses indicate that the EphA2 ICR exists as a dynamic configurational ensemble in solution. Rather than forming stable contacts and domain orientations, the kinase and SAM domains interact transiently, interconverting between a few cluster centers, with weak self-association of domains. This is reminiscent of the picture we derived of the EphA2 – SHIP2 SAM:SAM heterodimer interaction experimentally (Lee et al., 2012) and computationally (Zhang & Buck, 2014; Zhang et al., 2016; Li et al., 2018), which indicated that the protein-protein complex is a dynamic one. Here we find that the same appears to be true for the larger ICR in terms of its domain-domain intra-protein but also especially interdomain interactions. Modest intra-protein domain interaction and inter-protein binding affinities (Kd’s in the low μM regime) are typically characteristic of binary interactions in cell signaling functions as the protein complexes need to come apart on the second-to minute biological timescale, rather than being very tightly bound together. Still a hierarchy of interactions and their synergy or opposition play important roles in molecular mechanisms which regulate cell signaling. Thus, removal of the SAM domain enhances dimerization in both MST and simulations, indicating that SAM domain modestly inhibits EphA2 ICR self-association. This contrasts with crystallography-based perspectives (see comments in Borthakur et al., 2014; Singh et al., 2018), but aligns with fluorescence studies of full-length receptors in cells, which support dynamic clustering (Pasquale, 2024a; Shi et al., 2023, 2017; Singh et al., 2017). Collectively, our findings support a model in which EphA2 functions as a dynamic regulatory module whose conformational landscape is shaped by transient domain interactions, ultimately further facilitated by ligand-dependent signaling and membrane-associated organization (Gianni et al., 2021; Sahoo et al., 2025; Sahoo & Buck, 2021). Further studies will be required to elucidate how these factors collectively regulate EphA2 activation and function.

### Phosphorylation and SAM domain effects on dimerization

In-cell studies suggest clustering of the whole receptor in membranes promotes phosphorylation and activation (Borthakur et al., 2014; Kwon et al., 2018; Zapata-Mercado et al., 2025). In contrast, our solution-state MST and cross-linking data indicate that phosphorylated EphA2 ICR displays reduced dimer affinity, favoring transient association. This suggests phosphorylation may weaken dimerization, potentially facilitating receptor degradation or endocytosis. While MST can be sensitive to indirect effects (Bell et al., 2011; Li et al., 2021; López-Méndez et al., 2021a, 2021b) -see suppl. materials and discussion-, cross-linking confirms substantial dimer/oligomer formation in both ICR and ICRΔSAM. The constructs expressed in *E. coli* were partially phosphorylated at key tyrosines (Y772, Y588, Y594), but not within the SAM domain (Y921, Y930 and Y960) – see results for possible explanations. Importantly, at least in solution, phosphorylation does not promote persistent kinase–kinase interactions or clustering. The latter may only occur in the 2D space of the membrane. However, it should also be noted that fluorescence-based measurements of oligomerization report on longer distances and do not capture interaction lifetimes (Leavesley & Rich, 2016), so a transient, open configuration remains compatible with receptor-protein interactions when the ICR is tethered to membranes. Our results support the idea that many EphA2 phosphorylation sites primarily recruit SH2-containing effectors (e.g., Grb7, Shc), rather than enforce stable kinase proximity. Once activated, a loosely associated state may facilitate access of kinases, phosphatases, and adaptor proteins. High-resolution approaches such as cryo-EM or cryo-ET will be required to define receptor spacing within ligand-stabilized clusters, networks, or potential phase-separated assemblies (Borthakur et al., 2014; Case et al., 2019; Shi et al., 2023).

### Transient SAM interactions and regulatory implications

Consistent with CG simulations, AF2M, AF3 predictions and NMR, we detected no homodimerization of the EphA2 or SHIP2 SAM by themselves. Although our SAM deletion is artificial, disease-associated mutations destabilizing the SAM domain (Park et al., 2012), suggest physiological relevance, especially in cancer or neurodegeneration, where proteolysis may generate cytoplasmic fragments. These findings support continuing efforts to develop direct or allosteric EphA2 inhibitors (Liu et al., 2012; Mercurio & Leone, 2016; Petty et al., 2012; Schwalbe et al., 2026; Vincenzi et al., 2025). Adding isolated SAM domains to ICR or ICRΔSAM is not an outlandish proposition especially if EphA2 and SHIP2 SAM domains can spend a considerable time when they are relatively detached from other domains in the full length protein, which likely enables them to interact with other EphA2 inter-molecularly. Such interactions, indeed, increased EphA2’s kinase activity, apparently by competing with intramolecular SAM–kinase contacts. NMR revealed subtle chemical shift changes, mainly near helix 2 of the EphA2 SAM domain, consistent with transient interactions, albeit using a non-canonical interaction surface on the side of the EphA2 SAM domain. Together with MST, these data suggest that intramolecular SAM–kinase interactions and interactions with added SAM domains are of comparable strength, explaining the observed enhancement of transphosphorylation. EphA2 SAM and SHIP2 SAM domains form heterodimers via canonical end-helix/mid-loop interfaces (Knight et al., 2011). However, their highly complementary charge distributions confer substantial orientational flexibility, as seen in NMR and MD studies (Lee et al., 2012; Shi et al., 2017; Zhang et al., 2016). When isolated SHIP2 SAM is added to the ICR, it engages partly canonically and modestly enhances EphA2 activity.

Given their strong electrostatics and propensity for encounter complexes (Li & Buck, 2019; Zhang et al., 2016), SHIP2 SAM domains may also transiently contact kinase surfaces distinct from those used by EphA2 SAM. Although weak, such interactions could modulate signaling via local conformational, entropy or allosteric effects (e.g. Li et al., 2022). In monomers, SAM–kinase interactions may regulate JM-KD configurations, in turn limiting dimerization, whereas in dimers, SAM may shift to stabilize inter-protein contacts. Membrane attachment may further amplify these effects by increasing local concentration, restricting orientation, or enabling lipid-mediated interactions with cholesterol or PIP2 (Singh et al., 2024; Sahoo et al., 2025).

### Limitations of AlphaFold2-Multimer/AlphaFold3 (AF2M/AF3) and insights from simulations

Comparison of CG simulations with AF2M/AF3 revealed clear limitations of structure prediction. AF2M and AF3 failed to reproduce known asymmetric kinase dimer topologies observed in EGFR and EphA2 crystal structures (Jura et al., 2009; Nowakowski et al., 2002; Zhang et al., 2006) (Fig. S5 and Fig. S12). Notably, although AF3 produced a higher average interface confidence score (ipTM = 0.57) than AF2M, this increase in confidence did not translate into more accurate kinase dimer models. Interestingly, AF2M assigned higher confidence to ICRΔSAM dimers (score 0.62) than to full-length ICR (0.24), consistent with SAM impeding stable residue contacts required for dimerization. In contrast, CG simulations consistently revealed SAM interactions with the juxtamembrane region in both monomers and dimers, with multiple SAM orientations stabilizing distinct configurational states. The apparent coevolution of SAM and JM domains in Eph receptors (Kwon et al., 2018), supports a regulatory role for these contacts. Multiple independent CG runs yielded reproducible clusters, indicating robustness, while interconversion between these states in all four replicas supports a dynamic conformational ensemble. However, CG models lack atomic resolution, limiting interpretation of specific side-chain or electrostatic interactions. Future work combining atomistic simulations, mutagenesis, and experimental mapping by NMR or cross-linking will be essential. Overall, the limited success of AF2M and AF3 in modeling EphA2 ICR dimers likely reflects the broader challenge of identifying physiological interfaces for transient, weakly interacting complexes -a challenge that extends even to crystallography (Schweke et al., 2023). These results support an emerging view of EphA2 as forming dynamic, protein assemblies composed of multiple weak and context-dependent interactions (Gianni et al., 2021). The concept of certain protein-protein complexes being dynamic entities is gaining traction in the molecular biophysics community (e.g. Zhang & Buck, 2017; Lee & Lee, 2025). While the computational models are not definitive, compared to AF2M/AF3, the CG-derived ensemble is most consistent with our experimental observations (MST, NMR, cross-linking, and kinase assays), supporting a dynamic, asymmetric solution-state organization of the EphA2 ICR.

## Supplemental materials

Method and Materials

Tables S1-S7

Figures S1-S12

## Acknowledgements

We thank Drs. Francisco Barrera, Dimitar Nikolov, Adam Smith, Konstantin S. Mineev as well as Dr. Bing-Cheng Wang and for discussion. Dr. Zhen-Lu Li, Ms. Fatima Javier Razelle, and Ms. Deanna Bowman carried out early work on a precursor to this project. The project is partially supported by NIH grants R01AG089561 and previously R01EY029169 support to M.B. and the group. We want to thank CWRU HPC for providing computing resources.

