## Supplemental Tables and Figures for "Tyrosine phosphorylation and the inhibitory C-terminal SAM domain regulate the EphA2 intracellular fragment in solution: A combined experimental and molecular modeling study"

#### Materials and Methods

##### Cloning, expression and purification of EphA2 ICR

EphA2 ICR Full Length (559-976) and EphA2 ICR-ΔSAM (897-903) cDNA gene fragments, both from *Homo sapiens,* were codon optimized for expression in *E. coli*. The Protein constructs were inserted into a pET28a vector and also consist of a N-terminal thioredoxin (TRX) tag linked via a TEV recognition site (ENLYFQG) before a $His_6$-EphA2 sequence. We also expressed the same two proteins but with their tyrosine kinase function inactivated via residue mutation of Asp 739 to Asn. Plasmids were overexpressed in *E. coli* strain BL21 (DE3), and positive colonies were selected from Luria Bertani (LB) plates grown overnight with 0.05 mg/ml kanamycin. Colonies were grown in LB broth with 0.05 mg/ml kanamycin at 37°C at 200 rpm until the absorbance $O.D._{600}$ was between 0.6-0.8 units/cm. Protein expression was induced using 1 mM IPTG and growth continued at 25$^{o}$C 160 rpm. The cells were harvested after 16 hours or when the $O.D._{600}$ value was over 1.5. Protein expression, and eventually purity, was confirmed using SDS-PAGE (sodium dodecyl sulfate polyacrylamide gel electrophoresis). The cells containing the expressed protein, were resuspended in lysis buffer (25 mM TRIS pH 7.5, 300 mM NaCl, 5 mM $MgCl_2$, 2 mM TCEP ((Tris(2-carboxyethyl)phosphine) with a protease inhibition cocktail (Roche, Ref 04693159001) to prevent activity of *E. coli* proteases, giving a final volume of 50 ml which was then incubated with 1 mg/ml Lysozyme and DNAse for 20 mins to ensure proper lysis before a brief sonication on ice. The lysate was then centrifuged at 14,000 rpm at 4 $^{o}$C for 30 min and the supernatant was loaded onto 5 ml of Ni-NTA resin (Marvelgent)) equilibrated with 25 ml lysis buffer and washed with the lysis buffer containing 60 mM imidazole. The protein was eluted with 10 ml of the same buffer but containing 250 mM imidazole. Finally, after treating the elution for 18 hours TEV protease (0.3-fold concentration of eluted protein) at 20$^{o}$C to cleave off N-term TRX tag, it was subjected to Size Exclusion Chromatography, SEC, using Superdex 75 (Cytiva) in the buffer used for all subsequent experiments (20 mM TRIS pH 7.2, 150 mM NaCl, 5 mM $MgCl_2$ 2 mM TCEP) to separate the EphA2 protein from the cleaved TRX fusion tag. Our purification protocol differed from those used by Stauffacer (Balasubramaniam et al., 2011; Zabell et al., 2006) in that we used TEV protease for fusion tag cleavage instead of Factor Xa in a final buffer of 20 mM TRIS pH 7.2, 150 mM NaCl, 5 mM $MgCl_2$ 2 mM TCEP. Purity of the final sample was confirmed using SDS-PAGE and LC-MS. The SAM domains of EphA2 and SHIP2 were prepared as described by us previously (Lee et al., 2012).

##### Isotope ($^{15}$N) labeling and NMR spectroscopy

For hetero-nuclear NMR, we prepared an isotope $^{15}$N- labeled form of the ICR, ICRΔSAM, together with the SAM domains of EphA2 as well as SHIP2 SAM. Initially, cells were cultured in unlabeled M9 minimal medium at 37$^{o}$C and 220 rpm until the $O.D._{600}$ reached 0.6-0.7. The cells were then gently centrifuged and resuspended in PBS buffer to remove all the original M9 medium and centrifuged at 4500 rpm once more at room temperature; the cell pellets were then suspended in $^{15}$N ammonium chloride [$^{15}NH_4Cl$ 99%, Cambridge Isotope Laboratories, Inc.] M9 media, and induction was done at $O.D._{600}$ 0.5-0.6. Purification was done as described as above and for the SAM domains, as reported previously(Lee et al., 2012) . All NMR experiments were performed at 25$^{o}$C on Bruker Avance II 700, 800/900 MHz spectrometers equipped with a TCI-prodigy and TXI cryoprobes, respectively.

#### Coarse-grain (CG) molecular dynamics simulation

For modeling of the whole ICR of human EphA2 (res. 559-976), we chose the available crystal structure of inactive kinase domain (PDB ID: 4PDO) and the solution NMR structure of the SAM domain (PDB ID: 2KSO) from the Protein Databank (www.rcsb.org). We modelled the JM region, missing loops in the kinase domain and the linker region between the KD & SAM as a connecting loop or in the extended conformation of with the native sequence in PyMOL (The PyMOL Molecular Graphics System, Version 2.5. Schrödinger, LLC). ICRΔSAM (res. 559-903) was also modeled accordingly without the SAM domain, notably containing the full kinase-SAM domain linker.

The atomistic (AT) models of ICR/ΔSAM ICR monomers, modeled above, were transformed into a coarse-grained (CG) representation using the martinize2.py workflow module from the MARTINI 3 force field(Souza et al., 2021). Using the secondary structure assignments from DSSP(Kabsch & Sander, 1983), we employed an elastic network to enhance the stability of the secondary structure in the Kinase and SAM domains. We used default values of the force constant of 500 kJ/mol/nm$^{2}$ with the lower and upper elastic bond cut-off to 0.5 and 0.9 nm, respectively. CG simulations were performed using GROMACS version 2016.5.64 (Abraham et al., 2015) The pH of the systems was considered as 7.0, meaning that His are deprotonated (HSD?). All the simulations were run in the presence of regular MARTINI water and any net charge was neutralized and 0.15 M NaCl was added. To analyze the various possibilities for dimerization, the proteins were positioned 5.0 nm apart from each other. The systems were equilibrated for 500 ps. The long-range electrostatic interactions were used with a reaction type field having a cutoff value of 1.1 nm. We used potential-shift-verlet for the Lennard-Jones interactions with a value of 1.1 nm for the cutoff scheme, the V-rescale thermostat with a reference temperature of 310 K in combination with a Berendsen barostat with a coupling constant of 1.0 ps, compressibility of $3 \times 10^{-4}$ bar$^{-1}$, and a reference pressure of 1 bar. The integration time step was 20 fs. We followed the similar protocol for the monomer ICR simulation and those of ICR/ΔSAM ICR monomers with added EphA2 SAM or SHIP2 SAM. All the simulations were run in quadruplicate for 4 µs each. Trajectory analysis was conducted using the integrated modules within GROMACS. For additional analysis and comparisons, the CG structures that were extracted and converted back to an all-atomistic (AA) representation utilizing the Backward tool CG2AT2 (Vickery & Stansfeld, 2021; Wassenaar et al., 2014) .

#### Protein Complex Structure Prediction

For the prediction of dimer models within the same set of complexes and EphA2 or SHIP2 SAM binding to ICR/ICRΔSAM, Alphafold2 Multimer (Evans et al., 2022) and the webversion of Alphafold3 was employed, for the latter ATP and Mg were included and found correctly modeled into the kinase active site. For AF2M, model confidence was evaluated using the combined ranking confidence score (ipTM + pTM). For AF3, pTM and ipTM scores were reported separately, where pTM reflects confidence in the overall predicted structure and ipTM reflects confidence in the relative positioning of interacting subunits. Higher values indicate greater prediction confidence. The five top-ranked models were examined and showed no substantial differences in the overall domain arrangement or interaction interfaces; therefore, only the highest-ranked model was used for subsequent comparisons.

**NMR titration between SAM domains and ICR of EphA2**

An unlabeled ICR region of EphA2 was dephosphorylated using calf intestinal phosphatase at 1 unit for 25 µg of EphA2ICR for 4hrs at 25°C prepared in the NMR buffer (25mM TRIS pH 7.2, 150mM NaCl, 5mM $MgCl_2$, 2mM TCEP with 10% $D_2O$) at a final concentration of 120 µM. The titration point shown is at a 1:1 molar ratio of ICR or ICRΔSAM and $^{15}$N-labeled SAM domain of EphA2 or SHIP2. SAM residues that are broadened as a result of binding/ exchange processes are indicated using a cut-off of 1 standard deviation below the mean of all relative intensities. Notwithstanding allosteric effects, these changes suggest binding at these residues and their location was mapped on to the PDB : 2kso structure using the assignments reported(Lee et al., 2012).

The fractional error (i.e. combined errors in the $I_{bound}/I_{free}$, scaled by the value of this ratio), were estimated from propagation of the signal-to-noise ratio of the individual spectra using the following standard equation for error propagation:

$$\frac{\delta I_{bound}/I_{free}}{I_{bound}/I_{free}} = \sqrt{\left(\frac{\delta I_{free}}{I_{free}}\right)^2 + \left(\frac{\delta I_{bound}}{I_{bound}}\right)^2}$$

Where, $\delta I_{bound}/I_{free}$ is the calculated error in the $I_{bound}/I_{free}$ ratio, $I_{free}$ is intensity level of a peak in the free spectrum, $I_{bound}$ is intensity level of the same peak in bound spectrum, $\delta I_{free}$ is the baseline noise level of the free spectrum, and $\delta I_{bound}$ is the noise level of the bound spectrum.

Chemical shift perturbation, CSP upon titration were calculated using the following equation:

$$\Delta\delta av = \sqrt{[(\Delta\delta 1H)^2 + (\Delta\delta 15N/5)^2]}$$

Where, Δδav, Δδ1H, and Δδ15N are the changes in 1H and 15N chemical shift changes upon titration, respectively.

NMR is known as the “go-to” technique for detecting weak or transient interactions (Vinogradova & Qin, 2012) and there have been several similar examples of using a smaller assigned domain used to probe the interaction surfaces in a bigger protein (Goksoy et al., 2008). However, as we see here perturbations to signal intensities and chemical shifts can be very modest in the case of using either EphA2 or SHIP2 SAM domains as probes, consistent with the MD stimulations where interactions are transient and averaged over several contacts. Furthermore, the contacts themselves involve sidechains (incl. long chain sidechains, such as Arg, Lys and Glu) in solvated environments which only affect the proteins mainchain structure in a very small way (Lukhele et al., 2013). In the case of adding EphA2 SAM to the much larger ICR/ICRΔSAM in solution, a caveat to the NMR data is that the broadening of signal is relatively uniform across the SAM domain sequence. However, amide resonances in the helix-2 region and residues close to it, stand out as amide resonances which have increased intensity relative to the EphA2 SAM domain alone (unbound SAM domain). Longer T1 and especially T2 relaxation times are consistent with a movement of motions to shorter timescales but dynamics changes may be local either reflecting the influence of direct transient contacts or with allosteric effects. Remarkably, the helix-2 is involved as an interaction sites in our computational modeling and contact sites are overlapping the SAM – kinase domain contact regions seen in the CG simulations of the monomer, dimer and monomer + additional EphA2 SAM domain.

#### Western Blot Analyses

Western blot analyses of the elution fractions from the Ni-NTA resin column, EphA2 ICR, ICRΔSAM and ICR-kinase dead (D739N) were conducted to examine the level of phosphorylation. Samples were separated on 12% SDS-PAGE and then transferred onto a polyvinylidene difluoride (PVDF) membrane at 100 volts for 45 mins. Membranes were blocked with Everyblot Blocking Buffer (Biorad) for 10 minutes at room temperature following manufacturer's protocol. The membranes were then incubated with a 1: 1,000 dilution of EphA2 pY594 antibody (Bioss bs-3126R) or a 1:1000 dilution of pan Phospho-Tyrosine antibody (ABClonal AP0905) for 1 hour. The membranes were rinsed 5 times with TBS supplemented with 0.5% Tween 20, gentle shaking, with 10 min incubation period, rocking at room temperature between each wash. After washing, the membrane blots were developed with a 1: 1,000 dilution of Horseradish peroxidase (HRP) conjugated anti-rabbit secondary antibody (IgG (H+L) SA00001-2, Proteintech) for 1 hr at room temperature and were then rinsed 5 times with TBS supplemented with 0.05% Tween 20. For chemiluminescent detection, a 1:1 reaction of luminol reagent and peroxide solution (Biorad) was used. Membranes were visualized with Azure c600 (Azure Biosystems). For the trans-phosphorylation assay 10 µM kinase-dead EphA2 ICR or ICRΔSAM (substrate) was incubated, respectively, with 250nM wildtype EphA2 ICRΔSAM or EphA2 ICR (enzyme) at 22 $^{o}$C and reaction was quenched with 6X SDS-sample buffer at 0, 1, 2, 5, 10, 20, 30, 60 and 90 mins which were then subjected to Western blot using p-tyrosine, pY594 antibody as described above.

Quantification of WB: Western blot data was quantified using imageJ software. Each image was converted to greyscale and analyzed using the mean value of a defined "region of interest". The region of interest was defined in the software equal to the size of the largest band in a given row. A similarly sized region with no bands was used to quantify the mean value of background. The grey mean value of each lane is recorded and the background for the lane is subtracted to give the mean value related to the phosphorylated protein sample. Relative net protein values are calculated from the inverse of these mean values.

#### LC-MS and Phosphopeptide Analysis

For LC-MS analysis, bands from SDS-PAGE gels were cut out, dehydrated in acetonitrile, reduced with Dithiothreitol (DTT) and alkylated with iodoacetamide. The proteins were digested with trypsin and chymotrypsin overnight at room temperature. Peptides were extracted in two aliquots with 30 µL of 50% acetonitrile with 5% formic acid. The aliquots were combined, evaporated, and then resuspended in 30 µL 1% acetic acid prior to LC-MS analysis.LC-MS analysis utilized a ThermoScientific Fusion

Lumos mass spectrometry system. The HPLC column was a Dionex 15 cm x 75 μm id Acclaim Pepmap C18, 2 μm, 100 Å reversed-phase capillary chromatography column. 5 μL samples were injected and analyzed utilizing an LC gradient from 2 to 70% acetonitrile over 120 minutes. To analyze the percentage of phosphorylation, a second LC-MS/MS experiment was performed. This parallel reaction monitoring (PRM) experiment involves the fragmentation of specific ions over the entire course of the LC experiment. To estimate the level of phosphorylation, chromatographic peak areas of the phosphopeptides were normalized by the total peak areas of the phosphorylated and the unmodified form of the peptide to derive the extent (percentage) of phosphorylation.

Mass Spectrometry typically provides an estimate of the extent of phosphorylation at particular sites. Here, E. coli-expressed and purified EphA2 ICR and ICRΔSAM constructs were found to be partially phosphorylated (~50% occupancy) at several key tyrosine residues previously identified in mammalian systems and implicated in EphA2 kinase regulation, including activation loop Y772 and juxtamembrane residues Y588 and Y594(Fang et al., 2008; Pasquale, 2024). Interestingly, none of the three tyrosines within the SAM domain (Y921, Y930, and Y960) were detected as phosphorylated in our preparation, whereas another study showed phosphorylation at Y921 and Y930, also after expression in *E. Coli*. This suggests that phosphorylation of these residues may require additional cellular factors, alternative kinase activity, or may be influenced by local structural accessibility and solution conditions. However, our previous study demonstrated that the structural stability of the EphA2 SAM domain and its interaction with other SAM domains, including SHIP2 SAM, are not significantly affected by its phosphotyrosine status (Borthakur et al., 2014).

#### Kinase Activity Assays

EphA2 kinase activity was measured with a PhosphoSens Continuous Fluorescent Intensity Kinase Assay (AssayQuant Technologies # CSKS-AQT0104K) specifically designed for EphA2 kinases. Peptide concentrations for these assays ranged from 0.0-5.0 μM and the assay was executed according to manufacturer instructions in 50mM HEPES pH 7.4, 150 mM NaCl, 5 mM $MgCl_2$, 1mM EGTA and 0.5mM TCEP(Kolb et al., 2008). Data was collected every minute with a Tecan Spark plate reader (Tecan US, Inc) at 30 °C with an excitation wavelength of 360 nm and an emission wavelength of 485 nm. To analyze kinase assay data, the background fluorescence is subtracted from each time point to generate the relative fluorescence units (RFU). Corrected RFU data is plotted against time, and the initial reaction rates (RFU/min) are calculated from the slope of the linear portion (0 to12mins) of the RFU/time curve.

#### MicroScale Thermophoresis for Binding Affinity

Microscale thermophoresis (MST) on a Monolith NT.115 instrument (Nanotemper Technologies) was utilized to characterize domain-domain binding interactions. MST experiments were conducted in-solution enabling us to work in native-like conditions and to determine the dissociation constant, Kd. Binding of the fluorescently labeled molecule (see below) with the non-fluorescent molecule forms a complex and is subjected into a temperature gradient environment, induced by infrared laser. Depending on the size, charge, and hydration shell of the molecule, an altered diffusion of the bound state, compared to the unbound state, is observed. The thermophoresis signal is then plotted against the concentration of the ligand, generating a binding curve and the dissociation constant, Kd, is obtained assuming a 1:1 binding model (Asmari et al., 2018) .

Using the Monolith RED-tris-NTA kit (MO-L018, Nanotemper), we attached a fluorescently labeled dye molecule to the his-tag of the target protein. A solution of 200 nM protein and 100 nM dye was incubated for 30 minutes at room temperature, after which it is centrifuged for 10 minutes at 4°C, 13,000 g. The supernatant was removed and further diluted to a final concentration of 40 nM labeled protein. The target protein was serially diluted in PBS-T (0.05% Tween 20) for a sample volume of 5 µL, which was combined with 5 µL of labeled protein. Samples were incubated at room temperature for 5 minutes before being loaded into capillary tubes (MO-K025, Nanotemper). MST was conducted at 80% excitation power and medium MST power. All experiments were done in triplicate and analyzed using MO Affinity Analysis v2.2.6 (Nanotemper). All MST experiments were run as triplicates and standard errors were calculated accordingly.

Some degradation of the recombinant EphA2-ICR protein was observed during incubation, which may account for the additional band detected in the original blot. To independently confirm the observed affinity differences, we employed MST with labeling on lysine labeling, which occurs on a very different site compared to the N-terminal His-tag (MO-L011, Nanotemper). Consistent with our primary measurements, the wild-type EphA2 ICR bound its interaction partner with lower affinity (Kd= 5.3 ± 3.2 µM; Fig. S2A), whereas removal of the C-terminal SAM domain increased binding affinity approximately 2.5-fold (Kd = 2.1 ± 0.9 µM; Fig. S2B). The agreement between two independent labeling strategies demonstrates that the reduced affinity of the full-length ICR is an intrinsic property of the protein rather than a labeling artifact, providing strong evidence that the SAM domain negatively regulates binding through intramolecular conformational constraints.

It is unclear whether the MST signal changes report on dimerization/oligomerization – certainly not in persistent complexes since they are not detected by gel filtration. This situation is similar to that encountered in our work and that of others on the plexin receptor ICR where mutagenesis and crystallography are consistent with the formation of dimers/trimers, but such species are not detected by several other techniques (Bell et al., 2011; Li et al., 2021). It should be noted that MST has been established as a technique with good reproducibility over the years, but also shows sensitivity to effects which are not yet fully understood (López-Méndez et al., 2021b, 2021a), including the possibility of detecting transient interactions.

##### Cross-linking

Both Wildtype and kinase dead, D739N EphA2-ICR and EphA2-ΔSAM were prepared in 20 mM HEPES pH 7.8, 150 mM NaCl. Two different crosslinkers, namely DSSO, disuccinimidyl sulfoxide (ThermoScientific) and DC-4, 1,4-bis[4-[(2,5-dioxo-1-pyrrolidinyl)oxy]-4-oxobutyl]-1,4-diazoniabicyclo[2.2.2]octane, dibromide (Cayman Chemical company) were prepared in DMSO at a stock concentration of 10mM. Following the instruction from Thermo Scientific, crosslinking mix was prepared by mixing 10µM proteins in 25~50-fold excess of crosslinker in a total of 50µL reaction volume. The reaction mix was incubated at 25$^{o}$C for 45 mins and crosslinking reaction was quenched by incubating the crosslinking mix at 25ºC for 30 mins with 20 mM TRIS pH 7.8. Crosslinking was visualized using SDS_PAGE and western blot using Anti-EphA2 (Antibodies.com A99111). We note that in Figure S1 it is difficult to compare the wt and kd protein lanes in part because the former are more smeared out/broad, likely due to the presence of a range of phosphorylated states which affect the proteins' running in the gel.

Li Z-L, Müller-Greven J, Kim S, Tamagnone L, Buck M. Plexin-Bs enhance their GAP activity with a novel activation switch loop generating a cooperative enzyme.

Cellular and Molecular Life Sciences : CMLS. 2021;78(3):1101–12. https://doi.org/10.1007/s00018-020-03571-2

López-Méndez B, Baron B, Brautigam CA, Jowitt TA, Knauer SH, Uebel S, et al. Reproducibility and accuracy of microscale thermophoresis in the NanoTemper Monolith: a multi laboratory benchmark study. European Biophysics Journal : EBJ. 2021a;50(3–4):411–27. https://doi.org/10.1007/s00249-021-01532-6

López-Méndez B, Uebel S, Lundgren LP, Sedivy A. Microscale Thermophoresis and additional effects measured in NanoTemper Monolith instruments. European Biophysics Journal : EBJ. 2021b;50(3–4):653–60. https://doi.org/10.1007/s00249-021-01529-1

Lukhele S, Bah A, Lin H, Sonenberg N, Forman-Kay JD. Interaction of the eukaryotic initiation factor 4E with 4E-BP2 at a dynamic bipartite interface. Structure (London, England : 1993). 2013;21(12):2186–96. https://doi.org/10.1016/j.str.2013.08.030

Pasquale EB. Eph receptor signaling complexes in the plasma membrane. Trends in Biochemical Sciences. 2024;49(12):1079–96. https://doi.org/10.1016/j.tibs.2024.10.002

Souza PCT, Alessandri R, Barnoud J, Thallmair S, Faustino I, Grünewald F, et al. Martini 3: a general purpose force field for coarse-grained molecular dynamics. Nature Methods. 2021;18(4):382–8. https://doi.org/10.1038/s41592-021-01098-3

Vickery ON, Stansfeld PJ. CG2AT2: an Enhanced Fragment-Based Approach for Serial Multi-scale Molecular Dynamics Simulations. Journal of Chemical Theory and Computation. 2021;17(10):6472–82. https://doi.org/10.1021/acs.jctc.1c00295

Vinogradova O, Qin J. NMR as a unique tool in assessment and complex determination of weak protein-protein interactions. Topics in Current Chemistry. 2012;326:35–45. https://doi.org/10.1007/128_2011_216

Wassenaar TA, Pluhackova K, Böckmann RA, Marrink SJ, Tieleman DP. Going Backward: A Flexible Geometric Approach to Reverse Transformation from Coarse Grained to Atomistic Models. Journal of Chemical Theory and Computation. 2014;10(2):676–90. https://doi.org/10.1021/ct400617g

Zabell KM, Laurence JS, Kinch MS, Knapp DW, Stauffacher C V. Expression and purification of the intact cytoplasmic domain of the human ephrin receptor A2 tyrosine kinase in Escherichia coli. Protein Expression and Purification. 2006;47(1):210–6. https://doi.org/10.1016/j.pep.2005.10.031

#### Supplemental Tables

**Table S1.** Comparison of the intramolecular interactions between top clusters from the ICR monomer CG simulation.

**1$^{st}$ Cluster (ICR monomer)**

| | | | |
|---|---|---|---|
| **JM** | L582 | R946 | **SAM** |
| | Q581, S579 | D942 | |
| **KD** | T773 | R946 | **SAM** |
| | G776 | D943 | |
| | R792 | Q938 | |

**2$^{nd}$ Cluster (ICR monomer)**

| | | | |
|---|---|---|---|
| **KD** | R860, Q855 | T931 | **SAM** |
| | D888 | R907 | |

**Table S2.** Comparison of the intermolecular hydrogen bond interactions between the crystal structure and the 1st clusters from the ICR dimer, ICRΔSAM dimer CG simulations.

**1$^{st}$-Cluster (ICR dimer)**

| **Chain A** | | **Chain B** | |
|---|---|---|---|
| **KD1** | M727, T795, S796 | R946 | **SAM2** |
| | A731 | K945 | |
| | R858 | G948 | |
| | R861 | I947 | |
| | K728 | T940, D943 | |
| **KD1** | F834 | Y791 | **KD2** |
| | S844 | R858 | |
| | Q848, Q852 | K793, S790 | |
| | D879 | R860 | |
| | D886 | R858, S796 | |
| | R890 | G662 | |
| **SAM1** | E911 | K728 | **KD2** |
| | R907 | N734 | |
| **JM1** | D573 | R950 | **SAM2** |
| | V574 | K945 | |
| | Y575, F576 | Q955 | |
| | S577, K578 | R950 | |
| **KD1** | D832 | R566 | **JM2** |
| | K828 | Q565 | |
| | G833 | V574 | |

**1$^{st}$-Cluster (ICRΔSAM dimer)**

| **Chain A** | | **Chain B** | |
|---|---|---|---|
| **KD1** | N831, D832 | R677 | **KD2** |
| | E857, R860 | C612 | |
| | R835 | G668, T605 | |
| | R858 | S635, S636 | |
| | R860 | E607 | |
| | R890 | Q669 | |

**PDB ID: 7KJA**

| **Chain A** | | **Chain B** | |
|---|---|---|---|
| **SAM1** | D967 | K702, R705 | **KD2** |
| | K956 | E815 | |

**Table S3.** Comparison of the intermolecular interactions between the available crystal structures of EphA2 ICR homodimers.

**PDB ID: 5EK7**

| Chain A | | Chain B | |
|---|---|---|---|
| **KD1** | Q656 | H673 | **KD2** |
| | K649 | N674 | |
| | E654 | N697 | |
| | E663, M667 | V713 | |
| | M688, E626 | R721 | |
| | T653 | C753 | |
| | V681 | E809 | |
| | E607 | V810, T812 | |
| | E679 | T838, M840 | |
| | K686 | N674 | |
| | R657 | N697 | |
| | Q599 | D701, K702 | |
| | K603 | Y813 | |
| | K684 | E815 | |
| | K633, T614 | Q848 | |
| | S611 | M851 | |
| | A621 | I875 | |
| **JM1** | N598 | F703 | **KD2** |
| | D596 | F703, K707 | |
| | A600 | G709 | |
| | Y594 | E706 | |
| | H592 | E710 | |

**PDB ID: 1MQB**

| Chain A | | Chain B | |
|---|---|---|---|
| **KD1** | G709, D841 | H673 | **KD2** |
| | D708 | K728 | |
| | K882 | N750 | |

**PDB ID: 4PDO**

| Chain A | | Chain B | |
|---|---|---|---|
| **KD1** | G709 | D799, T795 | **KD2** |
| | L716 | E787 | |
| | V717 | S790 | |
| | G718 | Y791 | |
| | Y813 | K793 | |
| | R816 | V601 | |
| | W819, L821 | D659 | |
| | S822 | R657 | |
| | R705 | H737, S796 | |
| | S712 | Y791, R792, K793 | |
| | Y813 | H737 | |
| | E815 | Y735 | |
| | Y818 | G662, E663 | |
| | D832 | Y652 | |
| | F834 | T653 | |
| | M840 | T784 | |
| | D841 | W783 | |
| **KD1** | I779 | E595 | **JM2** |
| | N823 | D596 | |

**Table S4.** Comparison of the binding interface interactions between AF2M and AF3 predicted top models of ICR and ICRΔSAM dimers.

**AF2M prediction (ICR dimer)**

| Chain A | | Chain B | |
|---|---|---|---|
| **KD1** | H824 | K917 | **SAM2** |
| | E825 | R957 | |
| **KD1** | N831 | Y791 | **KD2** |
| | Y791 | N831 | |
| **SAM1** | Q919 | E769 | **KD2** |
| | R957 | E825 | |
| | K917 | H824 | |

**AF2M prediction (ICRΔSAM dimer)**

| Chain A | | Chain B | |
|---|---|---|---|
| **KD1** | K684 | E679 | **KD2** |
| | E679 | K684 | |

**AF3 prediction (ICR dimer)**

| Chain A | | Chain B | |
|---|---|---|---|
| **KD1** | R890 | D967 | **SAM2** |
| | T812 | R957 | |
| | D841 | K956 | |
| | D708 | H954, R957 | |
| **SAM1** | H954 | D708 | **KD2** |
| | K956 | D841 | |
| | D967 | R890 | |
| | R957 | E815, D708 | |

**AF3 prediction (ICRΔSAM dimer)**

| Chain A | | Chain B | |
|---|---|---|---|
| **KD1** | Q669 | E679 | **KD2** |
| | R657 | S683 | |
| | E679 | Q669 | |
| | S683 | R657 | |
| | N734 | S635 | |

**Table S5.** Comparison of the binding interface interactions between AF2M and AF3 predicted top models of ICR and ICRΔSAM with EphA2 SAM and SHIP2 SAM.

**AF2M predictions**

| EphA2 ICR | EphA2 SAM |
|---|---|
| E820 | R957 |
| D708 | Q955 |
| D841 | K956 |

| EphA2 ICR | Ship2 SAM |
|---|---|
| K828 | S1257 |
| K917 | D1224, E1226 |
| G953 | N1220 |
| H954 | D1223 |
| K956 | D1230, D1235 |
| R957 | D1223, G1221 |
| A829 | K1258 |
| E825 | S1257 |

| EphA2 ICRΔSAM | EphA2 SAM |
|---|---|
| L895 | S915 |
| R894 | Q968 |
| E820 | K917 |
| E815 | R957 |
| D841 | K956 |
| D708 | H954 |

| EphA2 ICRΔSAM | Ship2 SAM |
|---|---|
| K778 | N1220 |
| R782 | E1198, D1223 |
| N823 | D1223 |
| R792 | D1230 |
| K655 | E1234 |
| T898 | R1192 |

**Table S6.** Comparison of the binding interface interactions between CG simulations of ICR and ICRΔSAM with EphA2 SAM and SHIP2 SAM.

**CG simulations**

| EphA2 ICR | EphA2 SAM |
|---|---|
| D879 | K917 |
| D886 | K956 |
| K882 | E914 |
| S899 | D967 |
| M840 | Y960 |

| EphA2 ICR | Ship2 SAM |
|---|---|
| K586 | G1221, D1223 |
| R566 | D1224 |
| L585 | E1226 |
| K583 | D1230 |
| R858 | E1234, E1235 |
| K863 | E1234 |
| N732 | D1238 |
| K863 | E1237 |

| EphA2 ICR ΔSAM | EphA2 SAM |
|---|---|
| D886, S901 | K917 |
| E815 | K956 |
| F887 | R957 |
| T812 | Y960 |
| D841 | S961 |
| S712 | D967 |

| EphA2 ICR ΔSAM | Ship2 SAM |
|---|---|
| S796 | N1220 |
| R562, R566 | E1226 |
| R562 | S1229, D1230 |
| R858 | E1237 |
| K793 | E1238 |
| R858 | A1239 |

**Table S7.** Comparison of the binding interface interactions between AF3 predicted top models of ICR and ICR ΔSAM with EphA2 SAM and SHIP2 SAM.

**AF3 predictions**

| EphA2 ICR | EphA2 SAM |
|---|---|
| R890 | D967 |
| E815 | R957 |
| D708, T812 | R957 |
| D841 | K956 |

| EphA2 ICR | Ship2 SAM |
|---|---|
| R890 | L1256 |
| K917 | E1226, D1224 |
| G953 | N1220 |
| H954 | D1223 |
| K956 | D1230, D1235 |
| R957 | D1223, G1221 |

| EphA2 ICRΔSAM | EphA2 SAM |
|---|---|
| R890 | D967 |
| E815 | K917 |
| D708 | H954, R957 |
| D841 | K956 |

| EphA2 ICRΔSAM | Ship2 SAM |
|---|---|
| R816 | N1220 |
| N823 | G1221 |
| K778 | I1231 |
| R816 | E1238 |

#### Figure S1

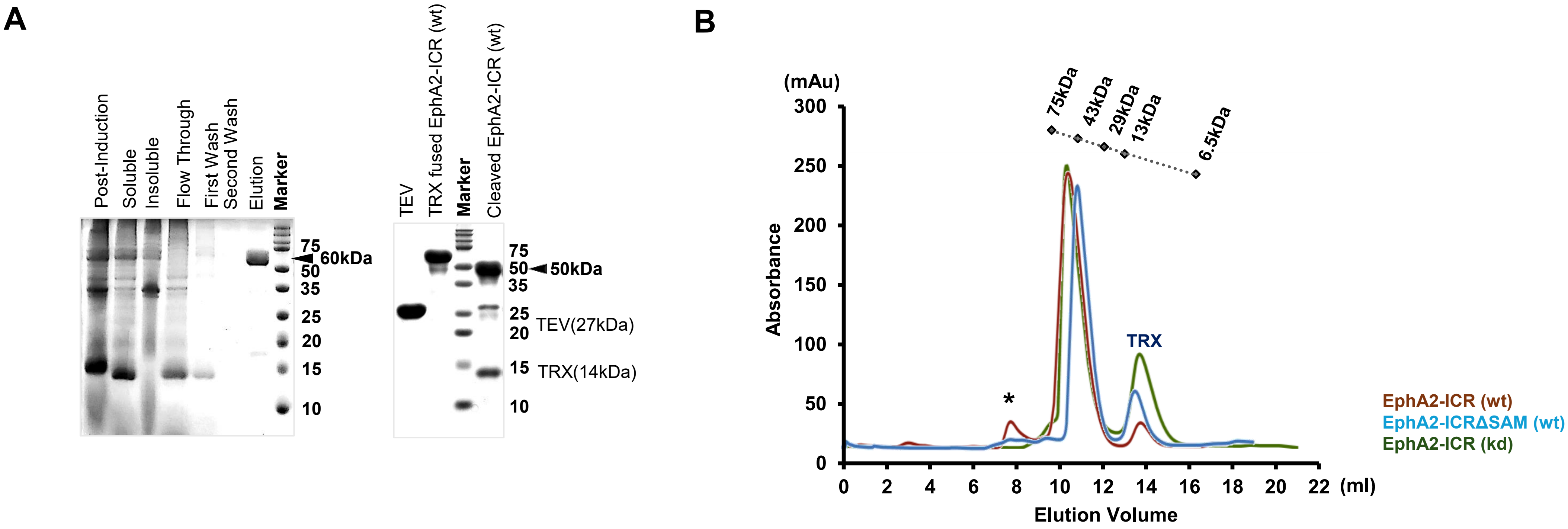


**Figure S1. Purification profile of the EphA2 intracellular region (ICR, ICRΔSAM)** (A) Affinity purification profile and protease cleavage of TRX fused EphA2 and, (B) Size exclusion chromatography using Superdex$^{TM}$ 75 10/300GL with standard calibration. Note the small peak at the column void volume, marked *, indicating some aggregation especially for the wt ICR, reduced for ICRΔSAM and essentially absent for kd ICR, consistent with a previous report on the stability/aggregation of the proteins

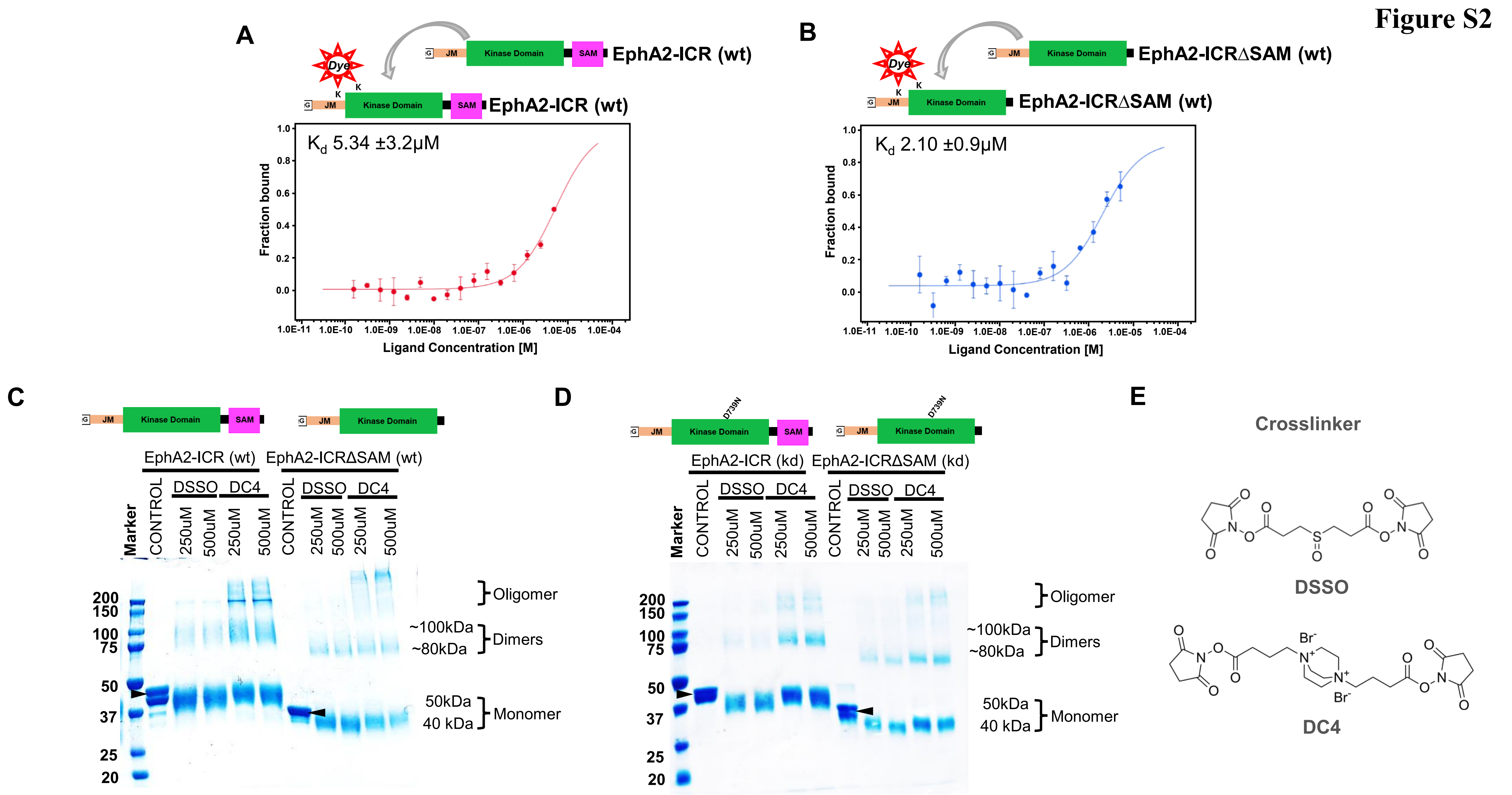


**Figure S2.** MST binding affinity using lysine labeling between (A) EphA2-ICR$^{wt}$ vs EphA2-ICR$^{wt}$, (B) EphA2-ICRΔSAM (wt) with EphA2-ICRΔSAM (wt). Cross-linking showing subtle difference between EphA2-ICR (wt) and EphA2-ICRΔSAM (wt) : (C) Wildtype, (D) Kinase-dead and (E) Chemical structure of crosslinker used.

### Figure S3

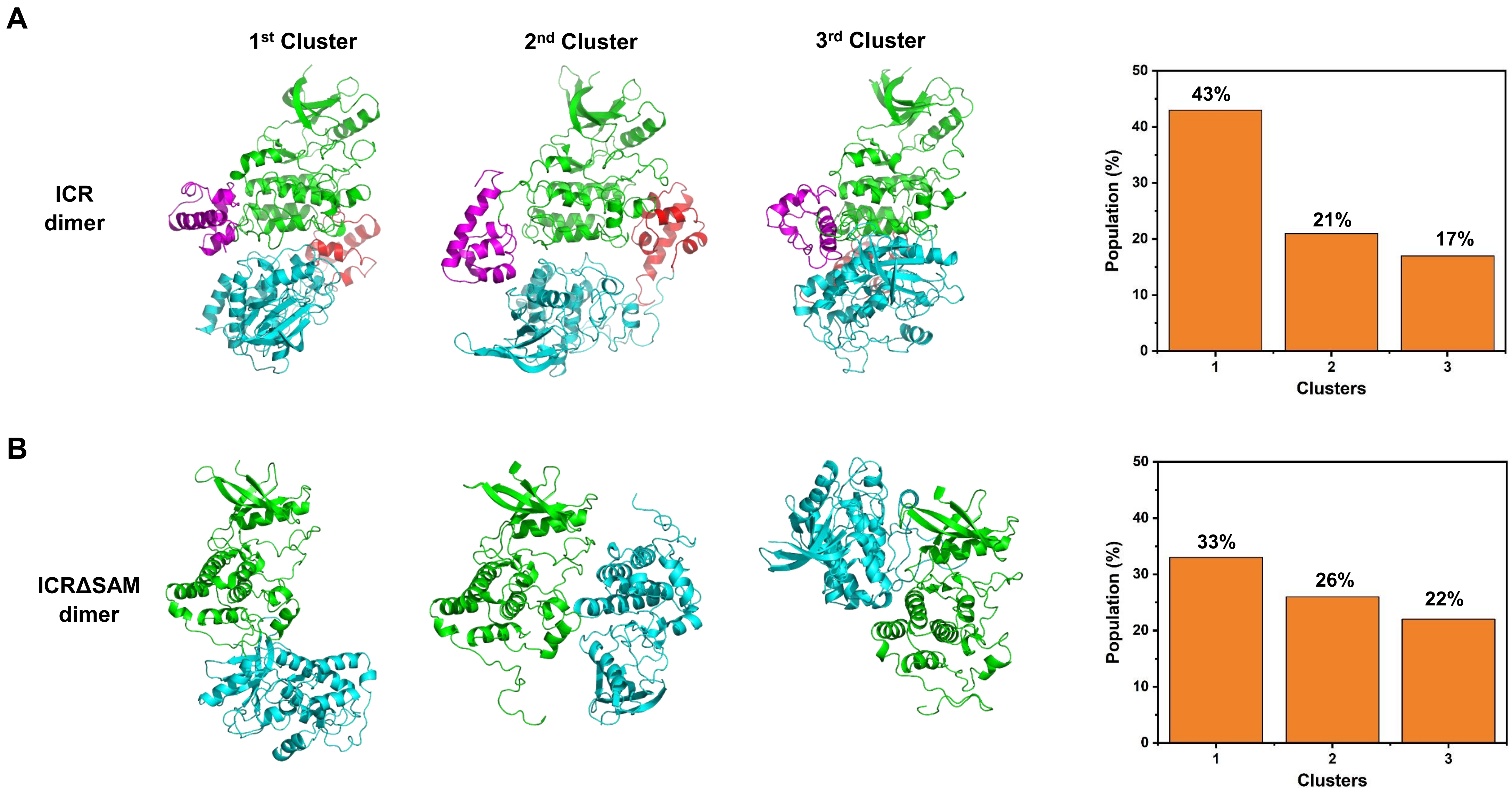


**Figure S3.** Comparison of the top three configurational clusters for ICR (upper panel) and ICRΔSAM (lower panel) dimer from the CG simulations. All the four replica simulations are clustered with a cutoff of 8Å. Coloring patterns for the cartoon representations are same as before.

### Figure S4

A

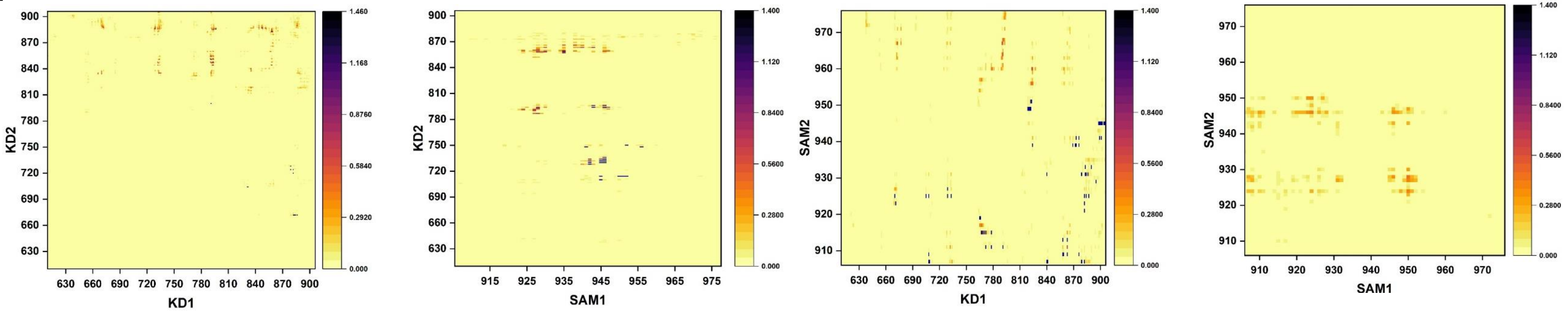


B

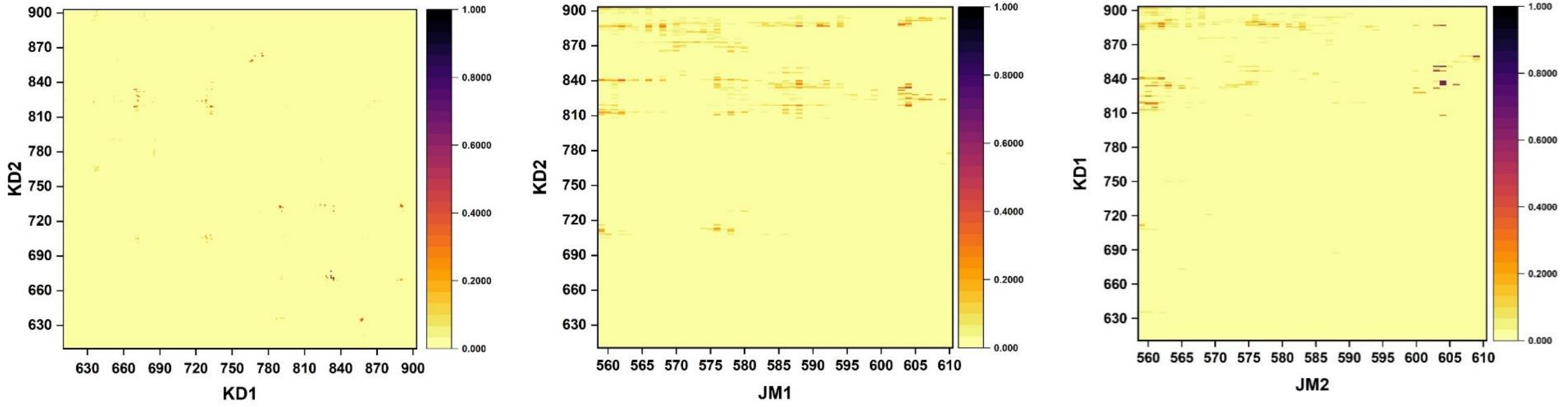


**Figure S4.** (A) Contact map of the dimerizing interface of ICR dimer over the course of the md simulation. Data from the last 1μs are merged from the 4 simulations, using a cut off 5Å for contacts. The color scale (yellow to brown to black) indicates the fractional occupation of contacts (0 to 1). Domains of chainA: KD1, SAM1; Domains of chainB: KD2, SAM2. (B) Contact map for ICRΔSAM. Domains of chainA: KD1, JM1; Domains of chainB: KD2, JM2.

### Figure S5

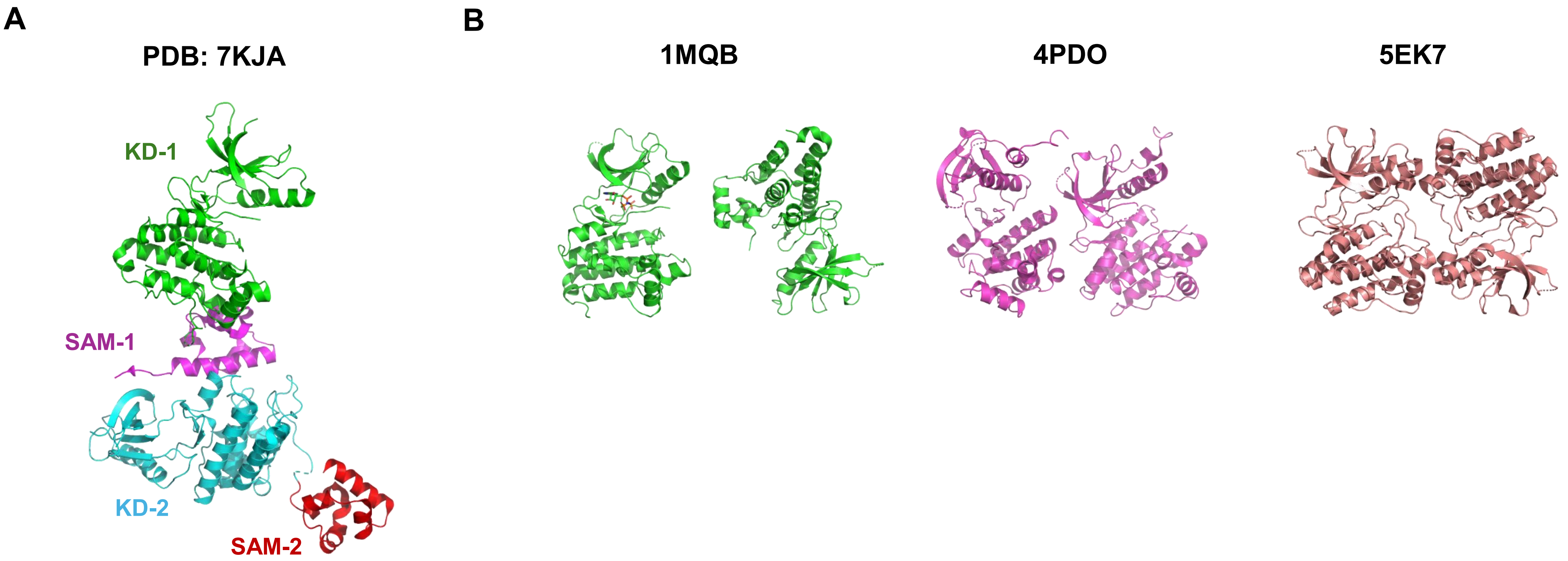


**Figure S5.** Comparison of the available crystal structures of EphA2 homodimers. (A) Cartoon representation of the wild-type intracellular region (ICR) of EphA2 containing both the kinase and SAM domains. The kinase domains are shown in green (chain A) and cyan (chain B), while the SAM domains are depicted in magenta (chain A) and red (chain B), matching the color scheme used in coarse-grained (CG) simulations for direct comparison. (B) Cartoon representations of other available structures of the wild-type EphA2 ICR containing only the kinase domains.

### Figure S6

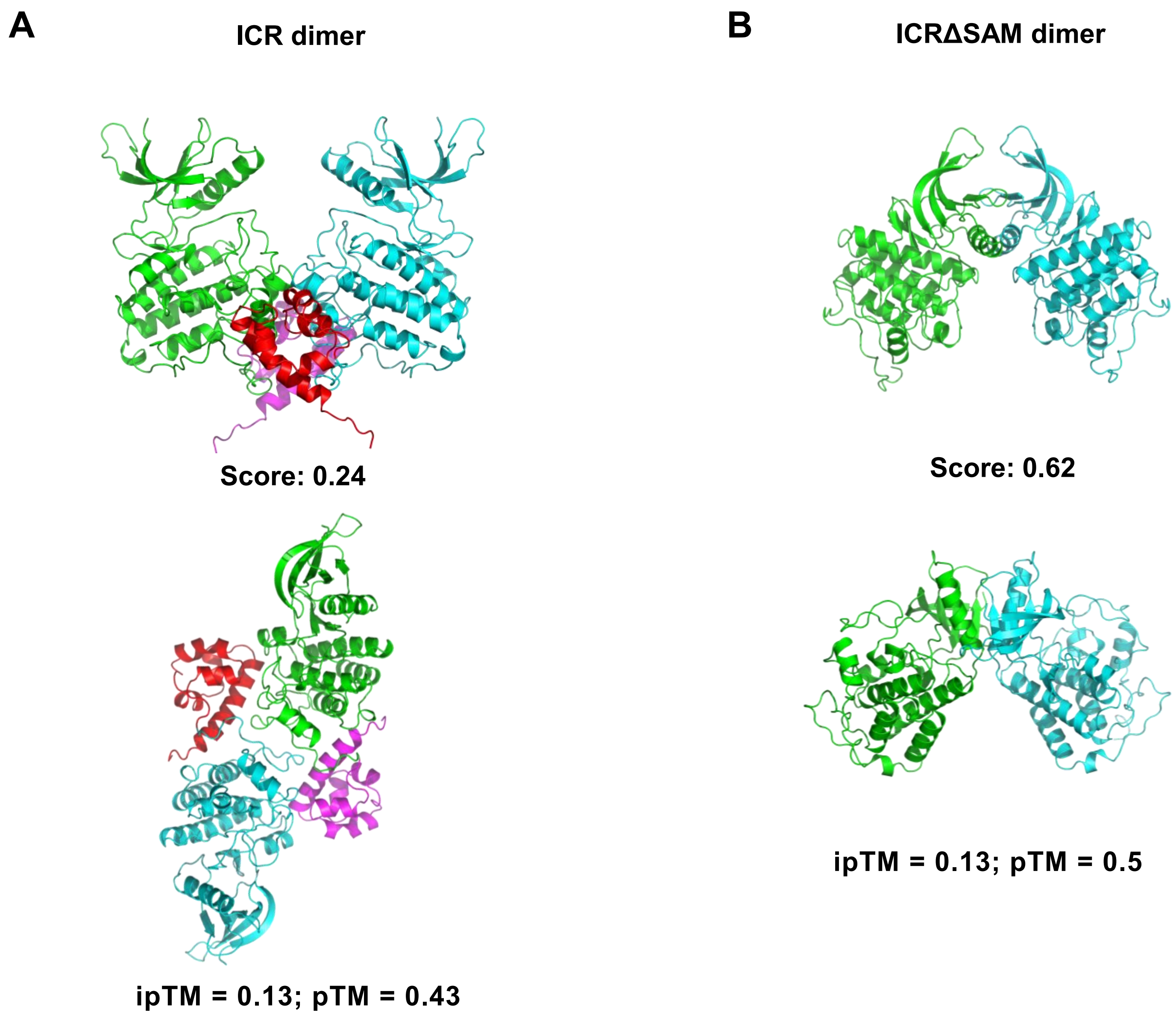


**Figure S6.** Comparison of AF2M (upper panel) and AF3 predictions (lower panel) of the homodimers of wt ICR and wt ICRΔSAM.

### Figure S7

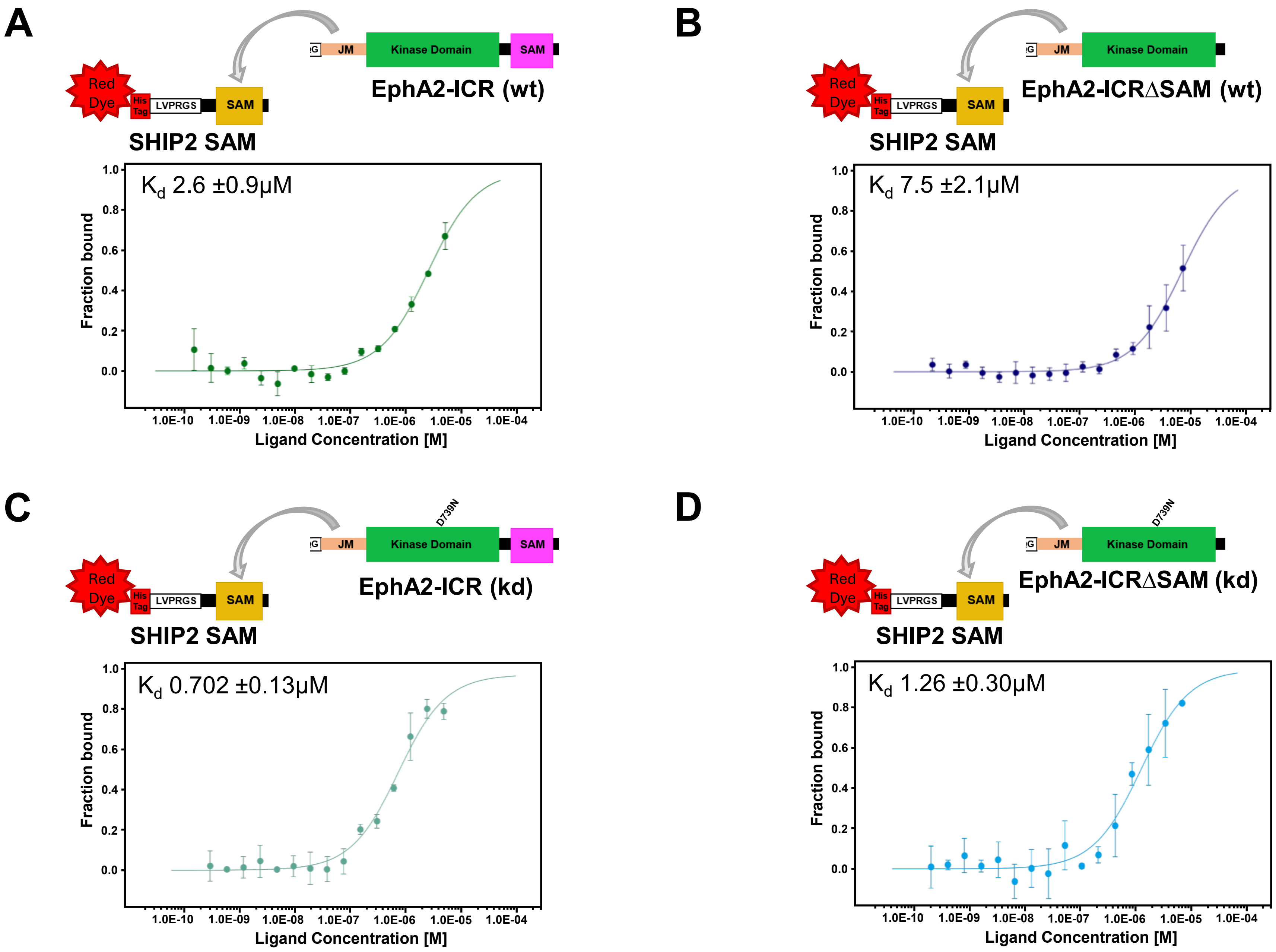


**Figure S7.** SHIP2 SAM domain bind to both ICR and ICRΔSAM regardless of phosphorylation state. MST binding affinity between (A) SHIP2 SAM vs wt ICR, (B) SHIP2 SAM Vs w ICRΔSAM , (C) SHIP2 SAM vs kd ICR and (D) SHIP2 SAM Vs kd ICRΔSAM.

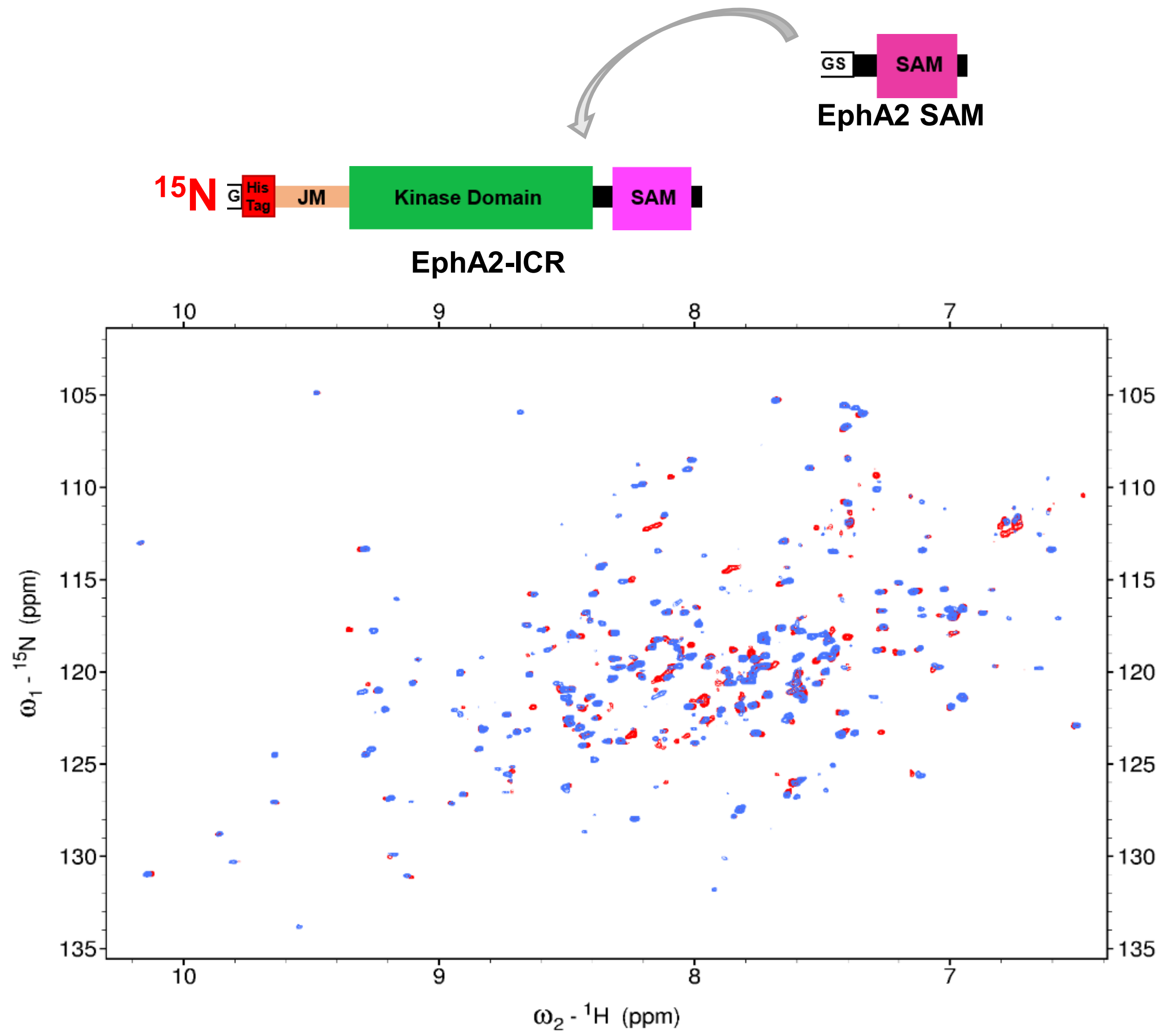


**Figure S8.** Chemical shifts seen in 15N labeled ICR upon binding of unlabeled EphA2 SAM. $^{1}$H-$^{15}$N TROSY-HSQC of 70μM ICR (de-phosphorylated) in 20mM TRIS pH 7.8; 150mM NaCl, 5mM $MgCl_2$ & 2mM TCEP with 2mM AMP-PNP at 303.1K (Red) and with unlabeled EphA2 SAM domain (Blue) at 1:1 molar ratio.

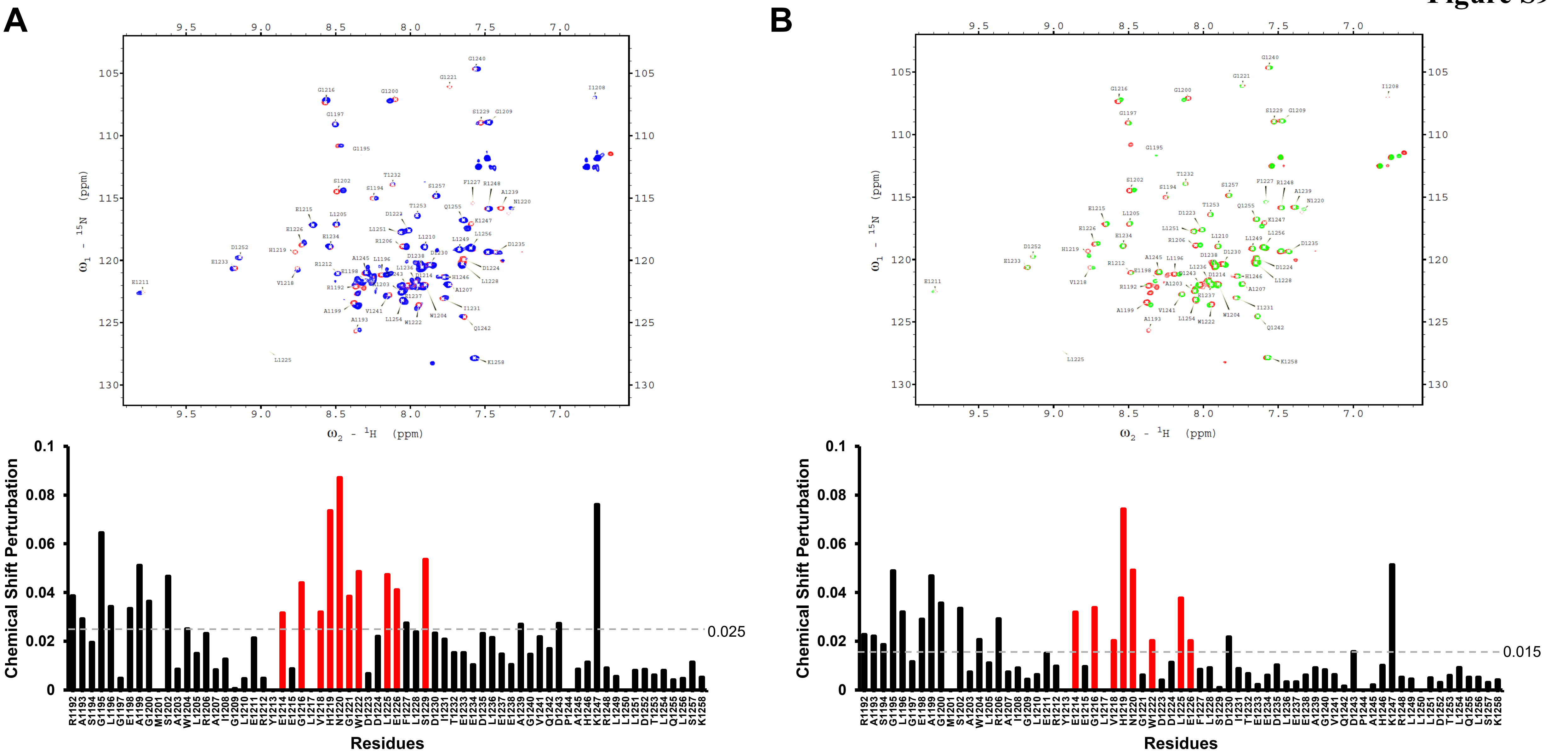


**Figure S9.** Chemical shift perturbation induced by interaction of ICR, ICRΔSAM on SHIP2 SAM domain. $^{1}H$-$^{15}N$ HSQC of SHIP2 SAM (Red) titrated with phosphorylated (A) ICR (Blue) and (B) ICRΔSAM (Green) at 1:0.3 molar ratio in 20mM TRIS pH 7.2; 150mM NaCl, 5mM $MgCl_2$ & 2mM TCEP at 298K at 700MHz. Chemical shift Perturbations were calculated using the equation Δδav = [(Δδ1H)2 + (Δδ15N/5)2]1/2, where Δδav, Δδ1H, and Δδ15N are the average, proton, and 15N chemical shift changes upon ICR (Blue)and ICRΔSAM (Green) titration, respectively.

### Figure S10

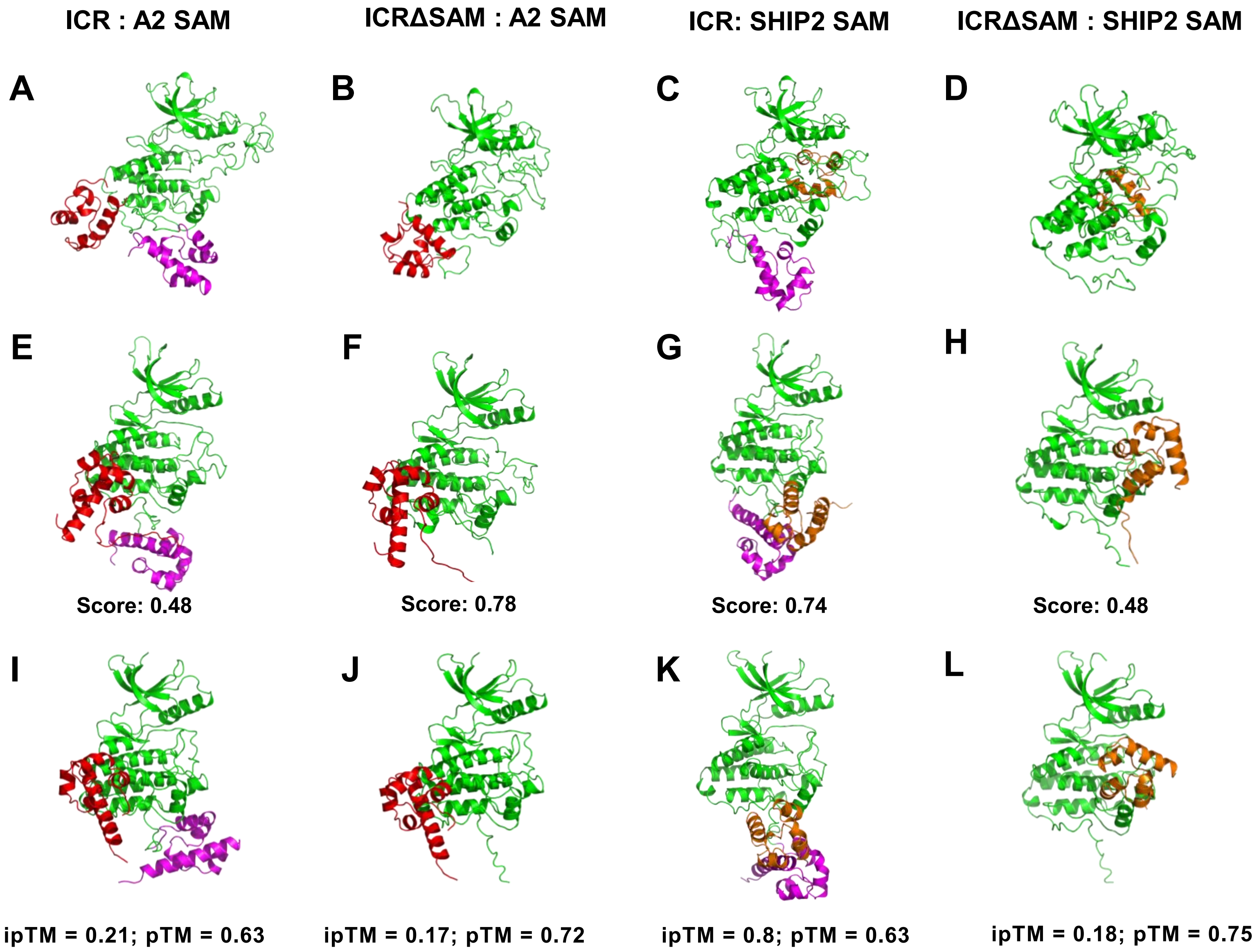


**Figure S10.** Comparison of top CG conformations (A-D), AF2M predictions (E-H) and AF3 predictions (I-L) for the protein-protein interaction between the ICR$^{wt}$ (KD in green and SAM in magenta) and ICRΔSAM (green) with EphA2 SAM (red) and SHIP2 SAM (orange).

Figure S11

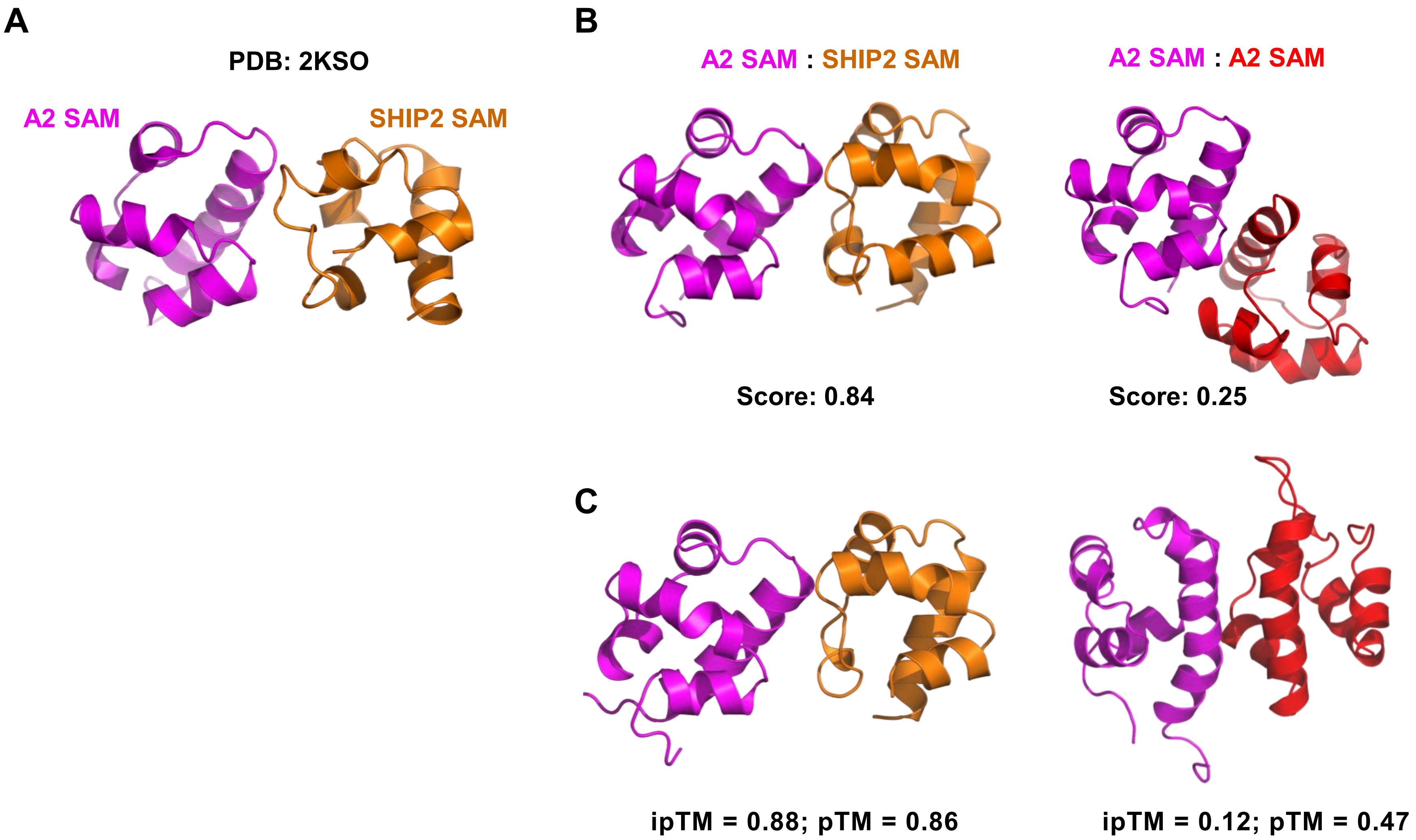


**Figure S11.** Comparison of NMR structure of the EphA2 SAM: SHIP2 SAM heterodimer (A) with AF2M predictions (B) and AF3 predictions (C) of the protein-protein interaction between the EphA2 SAM homodimer and for the EphA2 SAM with SHIP2 SAM heterodimer.

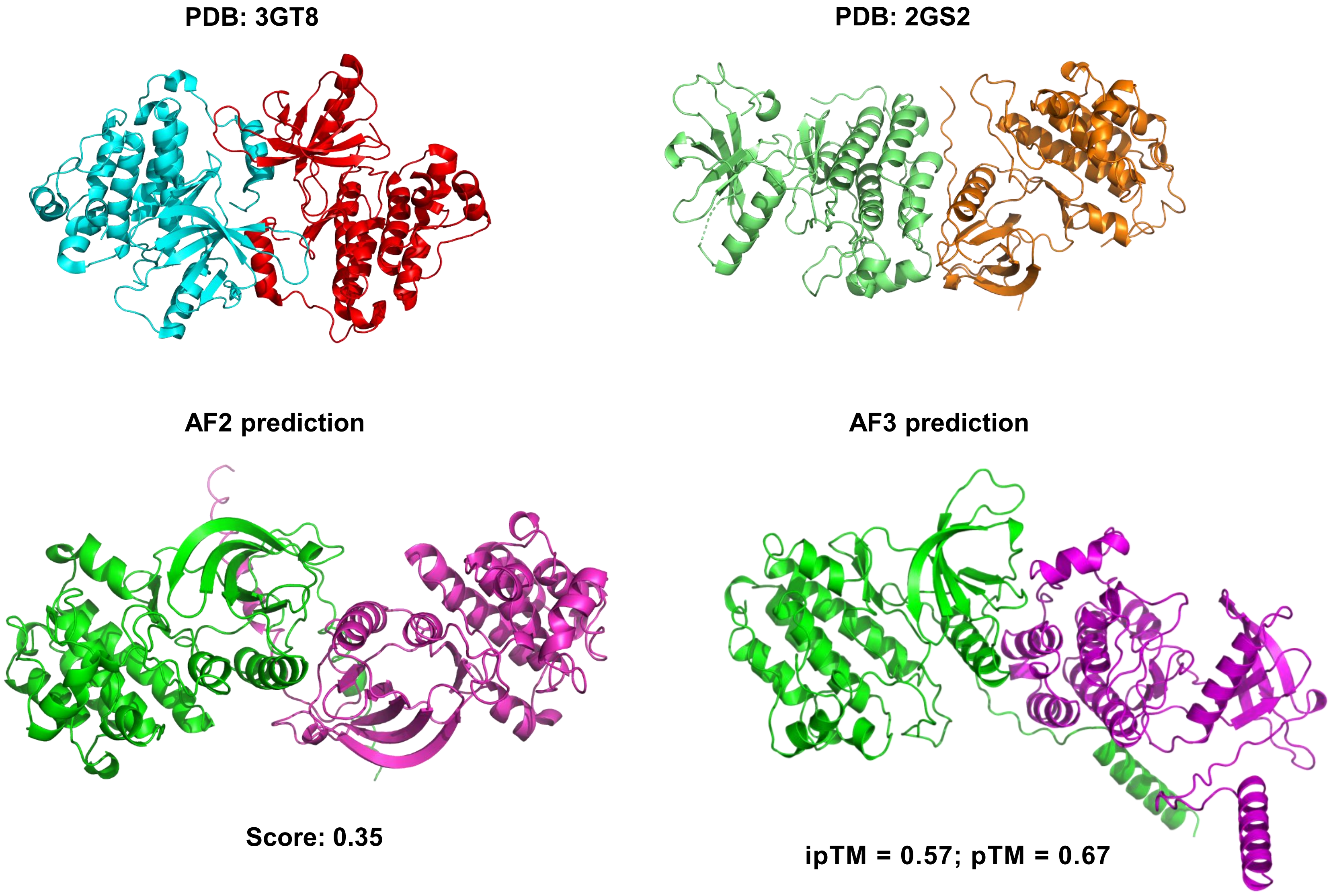


**Figure S12.** Comparison of the available crystal structures of EGFR dimer with the AF2M and AF3 predicted structures.
